# Nitrate restricts the expression of non-symbiotic leghemoglobin through inhibition of nodule inception protein in nodules of peanut (*Arachis hypogaea*)

**DOI:** 10.1101/2025.11.03.684091

**Authors:** Raju Kuiry, Swarup Roy Choudhury

## Abstract

An exquisite symbiotic relationship between legumes and rhizobia leads to the development of nitrogen-fixing special organelles known as nodules in nitrate-deficient environments, whereas a high level of nitrate in soil negatively regulates the pleiotropic phases of root nodule symbiosis (RNS), including rhizobial infection, nodule organogenesis and leghemoglobin synthesis. Here, we identified a special group of nodule-specific non-symbiotic leghemoglobin genes (*AhLghs*) in the crack entry legume peanut; however, their functional role and transcriptional regulation remain enigmatic. A comparative transcriptomic analysis revealed that the downregulation of nodule inception (*AhNIN*) and non-symbiotic leghemoglobin (*AhLghs*) genes played a pivotal role in the nitrate-mediated inhibition of nodulation in peanut. The knockdown of *AhLghs* and overexpression of *AhLgh1* resulted in lower and higher leghemoglobin content, respectively, corroborating their roles as positive regulators of nitrogen fixation in peanut. On the other hand, knockdown of *AhNINs* not only inhibited root nodulation but also decreased leghemoglobin content in peanut. Further, the DNA-affinity purification sequencing (DAP-Seq) analysis identified various nodulation genes, including *AhLghs*, as targets of AhNINs. After validating DNA-protein interaction by EMSA, the transactivation assay revealed that AhNINs can positively regulate *AhLgh1* after binding to the NIN RESPONSIVE CIS ELEMENT (NRCE) of its promoter. Our work bridges a critical gap in understanding how nitrate influences non-symbiotic leghemoglobin expression by targeting rhizobia-induced *NINs* in peanut, and offers a potential model suggesting that the nitrate-NIN-Lgh module might represent a key evolutionary event in fine-tuning root nodulation.

## Introduction

The uptake of nutrients from the environment is very crucial for plant growth and global crop productivity (Tilman et al., 2011; Mueller et al., 2012; Yadav et al., 2021; Rizzo et al., 2024). Nitrogen, a key component of nucleic acids, proteins, peptides, chlorophyll, and numerous secondary metabolites, is one of the essential nutrients in the soil to ensure healthy crops in agricultural fields. Although an enormous amount of nitrogen is available in the atmosphere, the availability of nitrogen in farming lands is often deficient worldwide. Therefore, cheaper synthetic nitrate fertilizers that predominantly contain nitrogen in the form of nitrate, ammonia or urea are extensively used to preserve soil fertility. The long-term excessive application of synthetic nitrogen fertilizers negatively affects soil fertility, crop yield and the function of the microbial community (Dai et al., 2018; Xu et al. 2020; Zhang et al., 2024). Thus, biological nitrogen fixation by root nodule symbiosis (RNS) is one of the best solutions for upholding sustainable agriculture (Oldroyd et al., 2011; Rogers and Oldroyd 2014; Udvardi and Poole 2013). But the RNS, which directly or indirectly promotes leguminous plant growth and development, is severely affected by the presence of excessive nitrogen fertilizers in soil (Streeter 1985; Van Noorden et al., 2016)

Unlike other plants, leguminous plants can fix atmospheric nitrogen (N_2_) by forming a symbiotic relationship with rhizobia, which they harbour in their root nodules (Oldroyd et al., 2011). Upon this interaction, plants deliver photoassimilates to the nitrogen-fixing rhizobia for their survival, and in turn, rhizobia provide fixed forms of nitrogen, i.e. ammonia, to their host (Lodwig and Poole 2003; Udvardi and Poole 2013; Battenberg and Hayashi 2022). Thus, legume plants can adapt effortlessly in nitrogen-deficient conditions where non-legume plants struggle for their survival. During RNS, the host plant releases flavonoids to induce the nod operon in soil-living rhizobia, resulting in the secretion of a lipo-chitin oligosaccharide, called Nod factors (NFs) (Oldroyd et al., 2011; Dong and Song 2020; Ghantasala and Roy Choudhury 2022). Subsequently, these factors are perceived by the Nod Factor Receptors (NFRs) present in the root hair epidermis to activate the symbiosis signaling pathway. This signaling cascade enables the onset of rhizobial infection and nodule organogenesis to house the incoming rhizobia. After colonisation in symbiosomes of nodules, rhizobia fix nitrogen with the help of the oxygen-sensitive nitrogenase enzyme complex, a central unit of nitrogen fixation. The inner cortex of the nodule acts as a diffusion barrier to limit the flux of oxygen to uphold a microaerobic condition suitable for the activity of nitrogenase (Ott et al., 2005; Avenhaus et al., 2016). Together with the inner cortex, legume plants deploy leghemoglobin, which is nearly 40% of total nodule proteins, to cope with the O_2_ paradox (Jiang et al., 2021).

Leghemoglobin that enables oxygen diffusion to the bacteroids and produces red colour in nodules is made up of a heme bound to an iron and a globin polypeptide chain (Avenhaus et al., 2016). Leghemoglobins are of two types in nodules, namely symbiotic leghemoglobins (Lbs) and non-symbiotic leghemoglobins or phytoglobins (Glbs) (Larrainzar et al., 2020). Although symbiotic leghemoglobins are expressed at millimolar concentrations in nodules, they exhibit canonical function in nodules. In contrast, non-symbiotic leghemoglobins are expressed at micromolar concentrations in different tissues of plants, including nodules, and are required to accomplish various developmental roles (Larrainzar et al., 2020; Villar et al., 2021). Three types of non-symbiotic leghemoglobins (Glbs), namely class 1, class 2 and class 3, show high, moderate and low oxygen affinity, respectively (Igamberdiev 2004; Smagghe et al., 2009) The structural adaptation of Lbs and Glbs, particularly within their heme pocket, is accountable for their diverse reactivity with oxygen; for example, Class 1 and 2 Glbs are hexacoordinate, but Lbs and Class 3 Glbs are pentacoordinate (Smagghe et al., 2009; Becana et al., 2020). Although hemes of classes 1 and 2 Glbs are also involved in the oxidation of NO, class 1 Glb mainly leads legume rhizobia symbiosis by NO homeostasis, whereas class 3 inhibits plant defence responses to promote symbiosis (Larrainzar et al., 2020; Villar et al., 2021). Studies on nodules of various models and crop legumes have identified multiple types of leghemoglobins, which are spatiotemporally regulated, and most of their individual physiological roles are not precisely annotated (Fuchsman and Appleby 1979; Uheda and Syono 1982; Uchiumi et al., 2002; Berger et al., 2020). The silencing of any class of Glb restricts vegetative growth and delays reproductive growth in *Lotus japonicus* by changes in the metabolome and hormone levels. Interestingly, the white nodule phenotype and the lower nitrogen fixing ability in nodules of the *lb123* mutant (devoid of three *LjLbs*) are only complemented by *LjLb2*, but not by *LjGlb*, suggesting a distinct functional diversity between symbiotic and non-symbiotic leghemoglobins (Villar et al., 2021).

Nitrate in soil not only restricts RNS but also inhibits leghemoglobin content in legumes (Becana et al., 1989). NIN-like proteins (NLPs) mediated signaling pathways are the primary cause of nitrate-induced control of root nodulation (Yan and Nambara 2023). The nitrate signaling activates NLPs through their N-terminal or amino-terminal regions, but the nodule inception (NIN), a member of a family of NIN-like proteins and a central regulator of root nodulation, is nitrate unresponsive due to its structural alteration in a region of the N-terminal domain (Suzuki et al., 2013). Except for this N-terminal region, the DNA-binding RWP-RK domain and protein-protein interacting PB1 domain are highly conserved between NLPs and NIN (Schauser et al., 2005; Konishi and Yanagisawa 2013). Comparable architecture of RWP-RK domain of NLPs and NIN facilitates them to bind a nitrate-responsive cis-element, NRE (Das et al., 2025). Predominantly, nitrate-responsive genes are transcriptionally activated by NLPs, whereas NIN antagonistically regulate nitrate-responsive genes to positively regulate root nodulation (Soyano et al., 2015). On the other hand, nitrate activates NLP1 for its translocation from the cytosol to the nucleus to physically interact with NIN, thereby inhibiting the transcriptional regulation of target genes, which are essentially required for root nodulation in *Medicago truncatula* (Lin et al., 2018). NITRATE UNRESPONSIVE SYMBIOSIS 1 (NRSYM1)/LjNLP4, a nitrate-responsive gene, dimerizes with NIN in *Lotus japonicus,* and the diverse DNA binding specificity of NLP4 and NIN modulates expression of genes required for nodulation (Nishida et al., 2021). A “double” version of the nitrate-responsive elements (NREs) existing in the promoter of symbiotic leghemoglobins is activated by NLPs in nodules of *Medicago truncatula* (Jiang et al., 2021). Although transcriptional regulation of several RNS genes is well established, it is mostly unclear how this intricate spatiotemporal expression of several leghemoglobins is transcriptionally monitored in nodules of different legumes.

The peanut (*Arachis hypogaea* L.), unlike other known model and crop legumes, allows crack entry for rhizobial invasion (Boogerd and Van Rossum 1997). Although symbiotic nitrogen-fixing capacity is high in peanut, the molecular signals involved in the establishment of nodules are still enigmatic. The availability of newer versions of the peanut genome enables us to identify gene sequences for further studies. To elucidate nitrate-mediated signaling cascades in peanut, we analysed symbiotic performance in response to nitrate in the preliminary part of this study. Phenotypic and transcriptome analysis revealed that nitrate treatment caused an inversion of leghemoglobin expression in nodules. We next identified non-symbiotic leghemoglobin categories and their functional relevance in peanut, which had not been explored properly in peanut. Further functional investigation by molecular approaches collectively revealed that NIN TFs guide non-symbiotic leghemoglobins in peanut nodules by a multifaceted stringent regulation of nitrate.

## Results

### Nitrate is a negative regulator of rhizobial infection and leghemoglobin expression in peanut

In response to rhizobia, *Arachis hypogaea* activates the expression of symbiotic genes for triggering nodule organogenesis to enhance nitrogen acquisition (Sinharoy et al., 2009). Although environmental nitrate promotes multiple adaptive responses favourable for plants, high nitrate levels in soil restrict pleiotropic phases of root nodule symbiosis (RNS) (Nishida and Suzaki 2018). When sufficient nitrogen is available in soil, nitrate-mediated signals halt symbiosis for the plant’s sake, as nodule development and rhizobial nitrogen fixation are energy-intensive processes (Nishida et al., 2020). To investigate the role of nitrate on nodule organogenesis and nitrogen fixation in peanut, 10-day-old peanut seedlings were treated with different concentrations of nitrate (10 mm and 20 mm) before and after rhizobia treatment (Supplementary Fig.S1). No significant difference in lateral root number was detected between control and 10-or 20 mm nitrate-treated plants, whereas a substantial increase in primary root length was observed in 20 mm nitrate-treated plants compared to the control as a result of nitrate-dependent root growth, perhaps via enhanced meristem activity and cytokinin signaling (Naulin et al., 2019) (Supplementary Fig.S2, A and B). Interestingly, a significant difference was observed in the nodule primordia and nodule number between the control (0 mm) and 20 mm nitrate treatment groups. About 80% of nodule primordia and 55% of nodule numbers were reduced in 20 mm nitrate-treated peanut plants (KR/nitrate plus rhizobia) compared to the control (R/rhizobia), whereas nodule numbers were identical in 10 mm nitrate-treated and control plants (Fig. 1, A to C). Several nodule sections of R and KR were performed to study the effect of nitrate on rhizobial colonization. Our observation clearly depicted that the rhizobial loading was extremely poor in 20 mm nitrate-treated plants compared to the control in peanut (Fig. 1D; Supplementary Fig. S3, A and B). One major difference between control and nitrate-treated nodules was nodule colour, which is mainly regulated by leghemoglobin (Lbs), a heme-containing protein essential for the protection of the nitrogenase enzyme (Larrainzar et al., 2020). Predominantly, non-nitrate-treated plants produced dark red nodule primordia and nodules, whereas light red to greenish and white nodules were detected in 20 mm nitrate-treated plants at both early and late nodulation stages (7 and 21 days post-infection or dpi) (Fig. 1E; Supplementary Fig.S4, A and B). About 4 to 5 times more white nodules were detected in 20 mm nitrate-treated plants, whereas rarely a few white nodules were observed in control plants, suggesting the involvement of nitrate in controlling the invasion of rhizobia and Lbs formation in peanut nodules (Supplementary Fig. S4C). Our quantitative measurement of Lbs content in control and 20 mm nitrate-treated plants also experimentally revealed that ∼60% reduction of Lbs in KR plants compared to R (Fig. 1F). To examine whether Lbs deficiencies lead to nitro-oxidative stress in peanut nodules, we detected the reactive oxygen species (ROS) in R and KR plants nodules. Our cytochemical staining (Nitrotetrazolium Blue Chloride/NBT and 3, 3, 9-Diaminobenzidine/DAB) exhibited the presence of an enormous amount of superoxide anion and hydrogen peroxide in KR nodules compared to R (Fig. 1, G and H). Further, a 4.5-fold more nitrate accumulation was quantified in KR compared to R roots, validating the uptake of more nitrate by KR roots (Supplementary Fig. S4D). Overall, these results indicated that nitrate accumulation in roots not only hinders rhizobial invasion and nodule formation, but also diminishes Lbs content in residual nodules, thereby increasing reactive oxygen species to damage the nitrogenase enzyme complex essential for nitrogen fixation.

**Figure 1.**
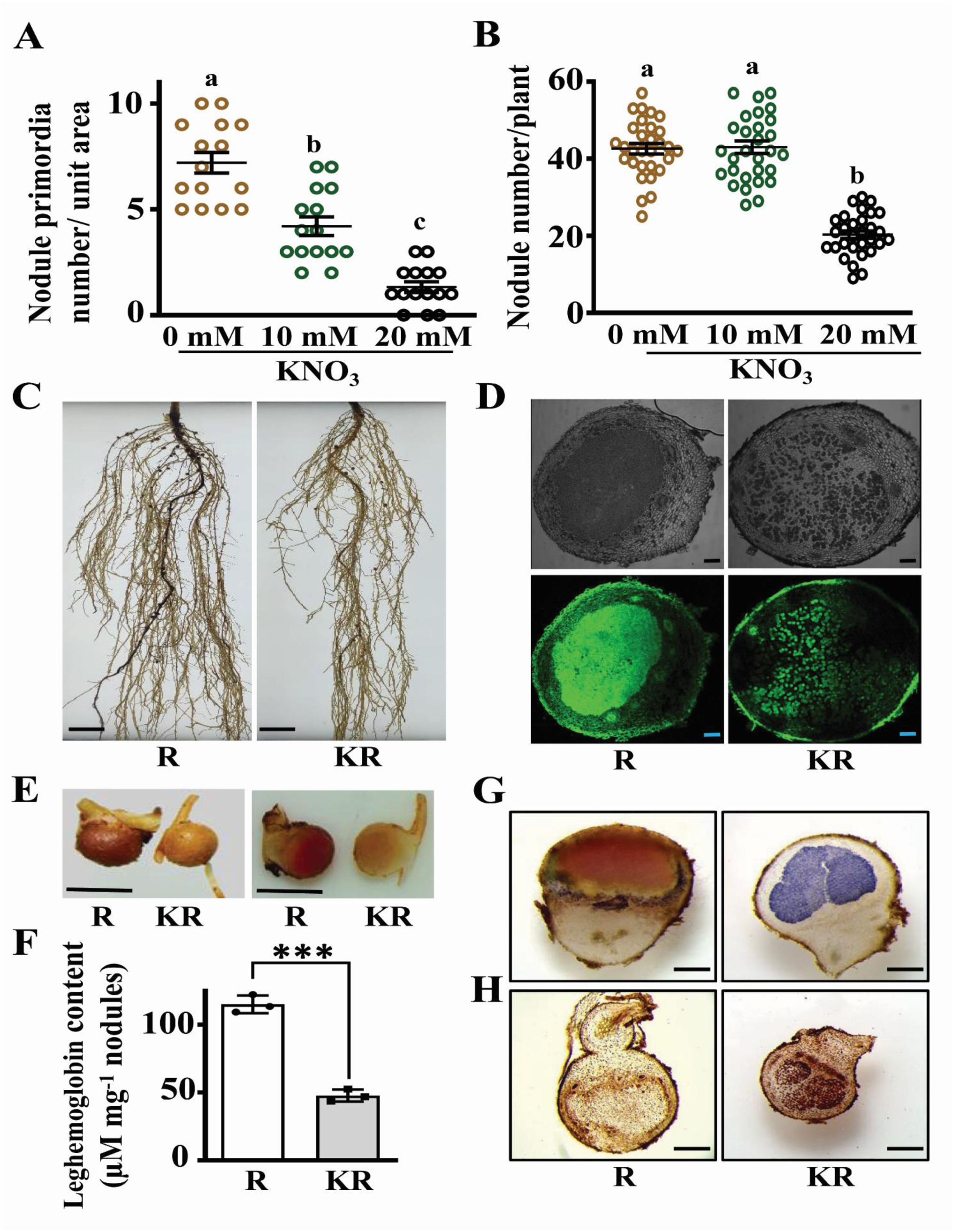
Nitrate is a negative regulator of rhizobial infection and leghemoglobin content in peanut. **A)** Peanut plants were grown in the absence and presence of different concentrations of nitrate (KNO_3_) and inoculated with rhizobia (*Bradyrhizobium sp. SEMIA 6144*). The nodule primordia number was counted per unit area (top 3 cm of each hairy root) of each root at 7-day post-inoculation (dpi). **B)** The nodule number per plant was estimated in the absence and presence of different concentrations of nitrate (KNO_3_) and rhizobia-treated plants at 21 dpi. Three biological replicates were used, and each replicate contained 5 to 7 roots and 10 to 12 plants for the estimation of nodule primordia and nodule number, respectively. Different letters indicate significant differences (Dunn’s multiple-comparisons test, P < 0.0001) between samples. Error bars represent ±SE. **C)** Representative image of 0-(R or rhizobia) and 20-mM nitrate (KNO_3_) (KR or nitrate plus rhizobia) treated roots at 21 dpi. Scale bars = 2 cm. **D)** Transverse sections of the nodules (40 µm thickness), obtained using a vibratome and stained with SYTO13, were observed under a confocal microscope (upper panel: brightfield and lower panel: SYTO13), respectively. Scale bars = 100 µm. **E)** Images of intact (left panel) and transverse sections (right panel) of nodules from 0-(R) and 20-mM nitrate (KNO_3_) (KR) treated roots represented a difference in colour. Scale bars = 2 mm. **F)** Estimation of leghemoglobin content in nodules of R and KR treated plants at 21 dpi. Asterisks indicate statistically significant differences (unpaired t test, ***P < 0.0001) between samples. **G)** Transverse sections of the nodules (80 µm thickness), obtained using a vibratome, were stained with NBT (nitroblue tetrazolium) to detect the superoxide radicals. Scale bars = 0.5 mm. **H)** Transverse sections of the nodules (80 µm thickness), obtained using a vibratome, were stained with DAB (3, 3’-diaminobenzidine) to detect the endogenous hydrogen peroxide. Scale bars = 0.5 mm.

### Transcriptomic analysis reveals nitrate-dependent downregulation of *leghemoglobin* and *NIN* genes

Leguminous plants reversibly regulate RNS depending on the amount of nitrate in the soil. To define potential regulators that affect multiple stages of nodule formation in the presence of nitrate, we performed a transcriptome analysis of peanut roots treated with 20 mm nitrate. After inoculating with rhizobia at 6 dpi, roots were collected from nitrate-treated (KR) and non-nitrate-treated (R) samples, and compared with uninoculated roots to gain further insight into how excessive nitrate impacts transcriptional regulation (Supplementary Fig. S5A). Volcano plot denoted the statistical significance distribution (-log10 p-value) versus magnitude of change as log2 fold change, where a diverse scattering of gene expression with the highest log2 fold change ranging from +10.0 (upregulated) to-10.0 (downregulated) was detected in both rhizobia treatment (R) and rhizobia plus nitrate treatment (KR) compared to control (Supplementary Fig. S5, B and C). This analysis identified a total of 5,813 genes that were upregulated 2.0-fold or more and 5,242 genes that were downregulated 2.0-fold or more in rhizobia vs control (R vs control), while 5,111 genes were upregulated and 6,468 genes were downregulated in rhizobia plus nitrate (KR) vs rhizobia (R) roots (Supplementary Fig. S5D). A complete summary of all Differentially Expressed Genes (DEGs) can be found in Supplementary Table S1. Interestingly, when we compared upregulated genes in R vs control and downregulated genes in KR vs R, it was observed that a total of 2,031 genes were upregulated in R vs control and were also downregulated in KR vs R (Fig. 2A; Supplementary Table S2). Gene Ontology (GO) enrichment analysis of up-regulated R vs control and down-regulated KR vs R revealed enrichment of biological processes related to protein phosphorylation, transmembrane transport, signal transduction and RNA modification, etc. and cellular components related to cytoplasm and mitochondria, etc. A categorization based on molecular function exhibited the most frequently identified process related to protein binding, followed by ATP binding and RNA binding (Supplementary Fig. S6, A to C; Supplementary Table S3), signifying an important role of nitrate in mediating these processes. To further examine the possible functions of differentially expressed common genes that were upregulated in R vs control and downregulated in KR vs R, we performed KEGG (Kyoto Encyclopedia of Genes and Genomes) analysis of those genes. Genes annotated as plant pathogen interaction, plant hormone signal transduction, MAPK signaling pathway, nitrogen metabolism, spliceosome, homologous recombination and GPI-anchor biosynthesis were significantly enriched, indicating that these molecular aspects were largely influenced when rhizobia inducible genes were inhibited by nitrate (Fig. 2B; Supplementary Table S4).

**Figure 2.**
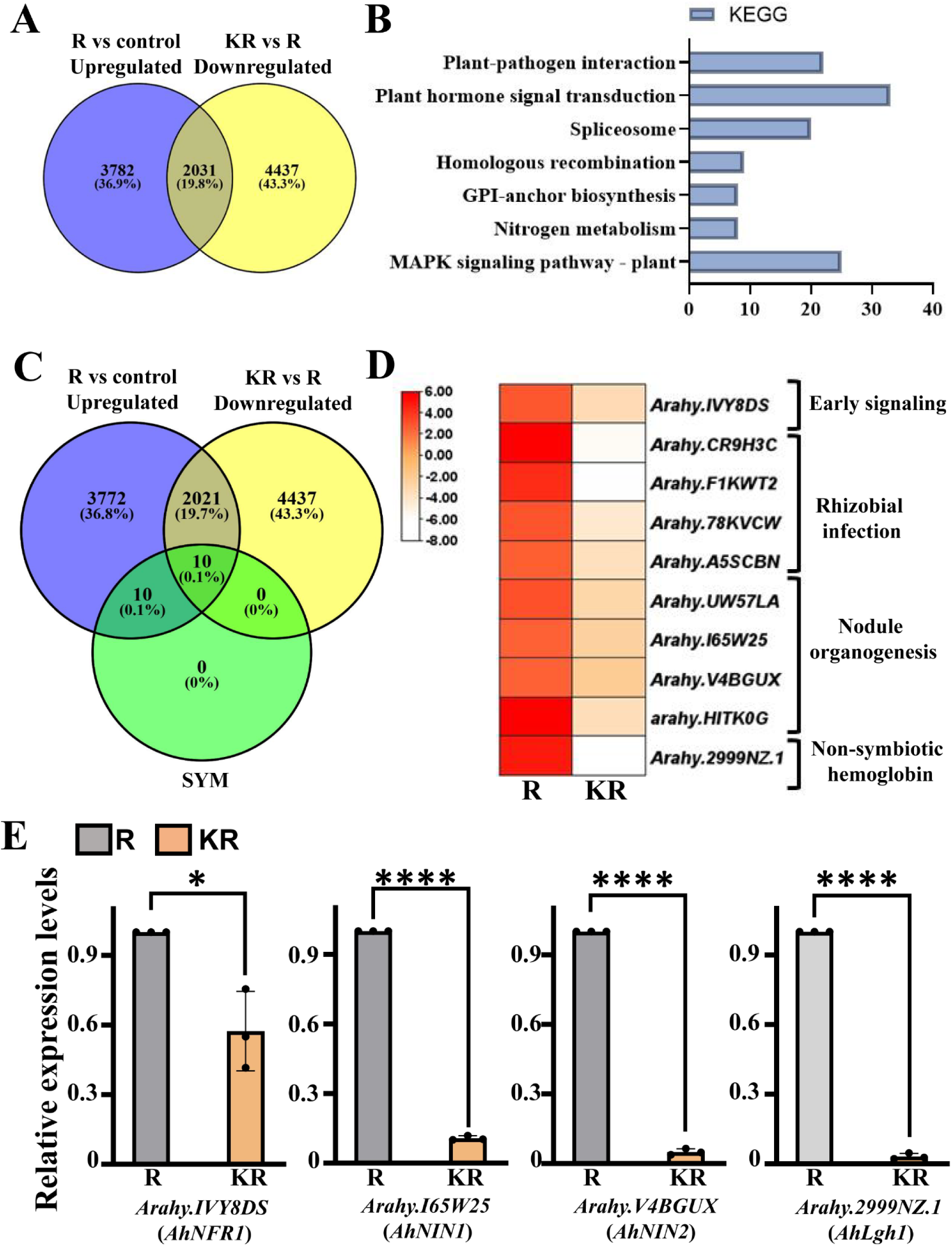
Transcriptomic analysis reveals the nitrate-mediated inhibition of *SYM* (symbiotic) genes in rhizobia-treated roots. **A)** A Venn diagram showed the overlap of DEGs among upregulated genes in rhizobia-treated roots compared to control (R vs control, blue) and downregulated genes in nitrate plus rhizobia-treated roots compared to rhizobia-treated roots (KR vs R, yellow) at 6 dpi. **B)** The bar graph showed the significant KEGG pathways identified from the common 2,031 genes between upregulated R vs control and downregulated KR vs R. The X-axis represented the number of DEGs/enzymes enriched in the top seven metabolic pathways. **C)** A Venn diagram showed the overlap of DEGs among upregulated genes in rhizobia-treated roots compared to control (R vs control, blue), downregulated genes in nitrate plus rhizobia-treated roots compared to rhizobia-treated roots (KR vs R, yellow), and SYM gene list derived from previous publications (green). **D)** The heat map exhibited expression levels of 10 common SYM genes without and with nitrate in R and KR roots at 6 dpi, derived from our RNA-seq data. Gene IDs and gene names were shown in **Supplementary Table S5**. The scale represented the log2FC values with upregulation in red and downregulation in white colour. **E)** The RT-qPCR validation was performed on selected genes from the transcriptome. Bar graphs denoted transcript levels of AhIVY8DS (*AhNFR1*), AhI65W25 (*AhNIN1*), AhV4BGUX (*AhNIN2*) and Ah2999NZ.1(*AhLgh1*) genes in nitrate plus rhizobia-treated roots (KR) compared to rhizobia-treated (R) roots at 6 dpi. The expression values were normalized against the peanut *Actin* gene expression. Expression in rhizobia-treated (R) roots was set at 1. Error bars represent the SE of the mean. Asterisks indicate statistically significant differences (unpaired t test, *P < 0.01, ****P < 0.0001) between samples.

To gain further insights into the symbiotic-related molecular candidates of rhizobia-induced and nitrate-suppressed, we compared 20 SYM (symbiotic)-related (previously published, Supplementary Table S5) with upregulated R vs control and downregulated KR vs R. We identified 10 common genes from SYM gene list (Fig. 2C; Supplementary Table S5), including AhIVY8DS (*AhNFR1*) (a marker gene for early symbiosis signaling), AhCR9H3C (*AhSymREM1a*), AhF1KWT2 (*AhSymREM1b*), Ah78KVCW (*AhNFH1*), AhA5SCBN (*AhROP6*) (marker genes for rhizobial infection), AhUW57LA (*AhLOG1*), AhI65W25 (*AhNIN1*), AhV4BGUX (*AhNIN2*), AhHITK0G (*AhNF-YA7*) (marker genes for early nodule organogenesis), Ah2999NZ.1 (*AhLgh1*) (a marker gene for leghemoglobin synthesis) (Fig. 2D) (Smit et al., 2007; Ke et al., 2012; Mortier et al., 2014; Cai et al., 2018; Liang et al., 2018; Schiessl et al., 2019; Laffont et al., 2020; Barker et al., 1988; Liu et al., 2021; Shrestha et al., 2021). In order to envisage the functional role of genes sharing similar expression dynamics, we performed co-expression analysis of rhizobia-induced but nitrate-suppressed genes. The STRING resultant interaction network, visualized and clustered in Cytoscape, discovered nine distinct functional modules. KEGG enrichment analysis showed that these clusters represent important biological processes such as plant-pathogen interaction, plant hormone signal transduction, lysine degradation, glycosylphosphatidylinositol (GPI)-anchor biosynthesis, nitrogen metabolism, purine and pyrimidine metabolism, starch and sucrose metabolism, amino sugar and nucleotide sugar metabolism and metabolic pathways that included mostly secondary metabolite biosynthesis, glycosaminoglycan degradation, inositol phosphate metabolism, arginine biosynthesis, other glycan degradation, galactose metabolism and glycerophospholipid metabolism (Supplementary Fig. S7; Supplementary Table S6). Within the network, *AhNIN* (Nodule inception) genes and their first neighbour interactors functioned as a hub gene to regulate all these biological processes, which shaped different clusters based on the KEGG pathway. Our analysis also revealed that one leghemoglobin gene (*AhLgh1*) and its first neighbour interactors acted as a strong hub. These co-expression clusters facilitated our understanding of the gene regulatory network in response to nitrate, and explained to us a possible mechanism that was adversely affected by high nitrate in the soil during RNS.

We identified 92 key transcription factors (TFs) involved in rhizobia-induced transcriptional regulation during nitrate suppression (Supplementary Fig. S8; Supplementary Table S7), where NIN-like transcription factors were mostly enriched. To confirm our transcriptome data, we studied the transcript abundance of both *AhNINs* and *AhLgh1* genes along with *AhNFR1* in our R and KR samples. Our gene expression profile exhibited that *AhNIN1*, *AhNIN2* and *AhLgh1* genes were severely affected in response to nitrate (Fig. 2E). Overall, our experimental results established that a substantial role of *AhNIN1*, *AhNIN2* and *AhLgh1* in nitrate-mediated signaling cascades of peanut during nodulation.

### Gene expression analysis exhibits nodule-specific non-symbiotic leghemoglobin genes downregulated by nitrate in peanut

To identify the complete repertoire of Lbs homologues, we queried the peanut database (https://www.peanutbase.org/, https://phytozome-next.jgi.doe.gov/info/Ahypogaea_v1_0) and subsequently studied their evolutionary relationships with Lbs of *Medicago truncatula* and *Lotus japonicus*. The resulting tree clearly segregated symbiotic leghemoglobins (Lbs) and non-symbiotic groups of globins (Glbs) of three genomes. Most of the Lbs of *M. truncatula* were closely related; only four members formed another cluster, which was closely associated with the GLb3 clade (Fig. 3A). The Lbs (Lb1–Lb3) of *L. japonicus* clustered adjacent to their *Medicago* counterparts but shaped a separate clade, indicating that both shared a common ancestor. Unexpectedly, we did not notice any homologue of symbiotic leghemoglobin in the peanut genome, but we detected one Glb1 (AhU2ETAV.1) and four Glb3 (Ah8TR5WT.1, Ah72Y1Y4.1, Ah LJ0T63.1 and AhYGZX15.1) in *A. hypogaea* with other Glb1s and GLb3s of *M. truncatula* and *L. japonicus*, respectively, which formed Glb1 and GLb3 specific clusters separately. Interestingly, seven non-symbiotic leghemoglobins (Ah2999NZ.1, AhH8G850.1, Ah9R3RK8.1, AhSU9WSI.1, AhZEE1Y7.1, AhC5DNVC.1, and AhSLEA2F.1) of *A. hypogaea* formed a distinct group from Lbs and Glbs of *M. truncatula* and *L. japonicus*. Out of seven, two sequences were partial. We considered the other five as a special group of non-symbiotic leghemoglobin genes (*AhLgh1* or *Arahy.2999NZ.1*, *AhLgh2* or *Arahy.H8G850.1, AhLgh3* or *Arahy.9R3RK8.1, AhLgh4 or Arahy.SU9WSI.1, and AhLgh5* or *Arahy.C5DNVC.1*) in the peanut genome (Fig. 3A). Our transcriptome data identified *AhLgh1* as a rhizobia-inducible gene, but inhibited after nitrate treatment (Fig. 2D). To further examine, we found a conserved N-terminal region among all *AhLgh* proteins, but their C-terminal regions were highly diverse (Supplementary Fig. S9A). There was a close similarity between AhLgh1 and AhLgh2 (98.67%), and AhLgh3 and AhLgh4 (96.37%), indicating the involvement of genome duplication events in the origin of these genes (Supplementary Fig. S9B). Chromosomal analysis exhibited localisation of Lgh1, Lgh4 and Lgh5 in chromosome 3, whereas Lgh2 and Lgh3 were localised in chromosome 13. A similar gene structure was observed in five AhLgh genes, and all contained 4 exons (Supplementary Fig. S10, A and B, (see details in Supplementary materials). Furthermore, the analysis of the AhLgh (AhLgh1-5) structures revealed three highly conserved motifs, but except AhGlb1, which were absent in AhGlb3-1, AhGlb3-2, AhGlb3-3 and AhGlb3-4 (Supplementary Fig. S11, A and B). Further, pairwise alignment and three-dimensional structural alignment using the UCSF ChimeraX MatchMaker tool revealed that AhGlb1 is structurally dissimilar from AhLgh1 (Supplementary Fig. S12, A and B, Supplementary materials). To decipher the structural conservation of leghemoglobin across legumes and non-legumes, the structure of a special group of non-symbiotic leghemoglobin gene (AhLgh1) of peanut was compared with its legume counterpart, i.e. leghemoglobin of *Medicago truncatula* (MtLb) and other non-legume hemoglobins (Hbs) like *Arabidopsis thaliana* (AtHb1) and *Oryza sativa* (OsHb1). Multiple sequence alignment and pairwise analysis revealed that AhLgh1 is highly similar to non-legume hemoglobins (AtHb1, OsHb1), in comparison to MtLb1 (Supplementary Fig. S13A). Similar to AtHb1 and OsHb1, AhLgh1 contained motif 1, which was absent in MtLb1 (Supplementary Fig. S13, B and C). Three-dimensional structural alignment using the UCSF ChimeraX MatchMaker tool further confirmed these findings (Supplementary Fig. 13, D and E). Pairwise alignment of AhLgh1 with AtHb1 and OsHb1 yielded low RMSD values and high alignment scores, indicating strong structural conservation. In contrast, alignments of AhLgh1 with MtLb1 exhibited higher RMSD values and low alignment scores, reflecting greater structural divergence (Supplementary Fig. 13, D and E; Supplementary Table S8). AhLgh1 shared a highly conserved globin fold, dominated by α-helices according to secondary structure prediction using DSSP (Define Secondary Structure of Proteins algorithm), which displayed more structural resemblance to non-legume hemoglobins (AtHb1, OsHb1) compared to MtLb1. Secondary structure prediction of Motif 1 also established the presence of a turn and random coil in MtLb1, which was absent in AhLgh1 and non-legume hemoglobins (AtHb1, OsHb1) (Supplementary Fig. 13, E and F).

**Figure 3.**
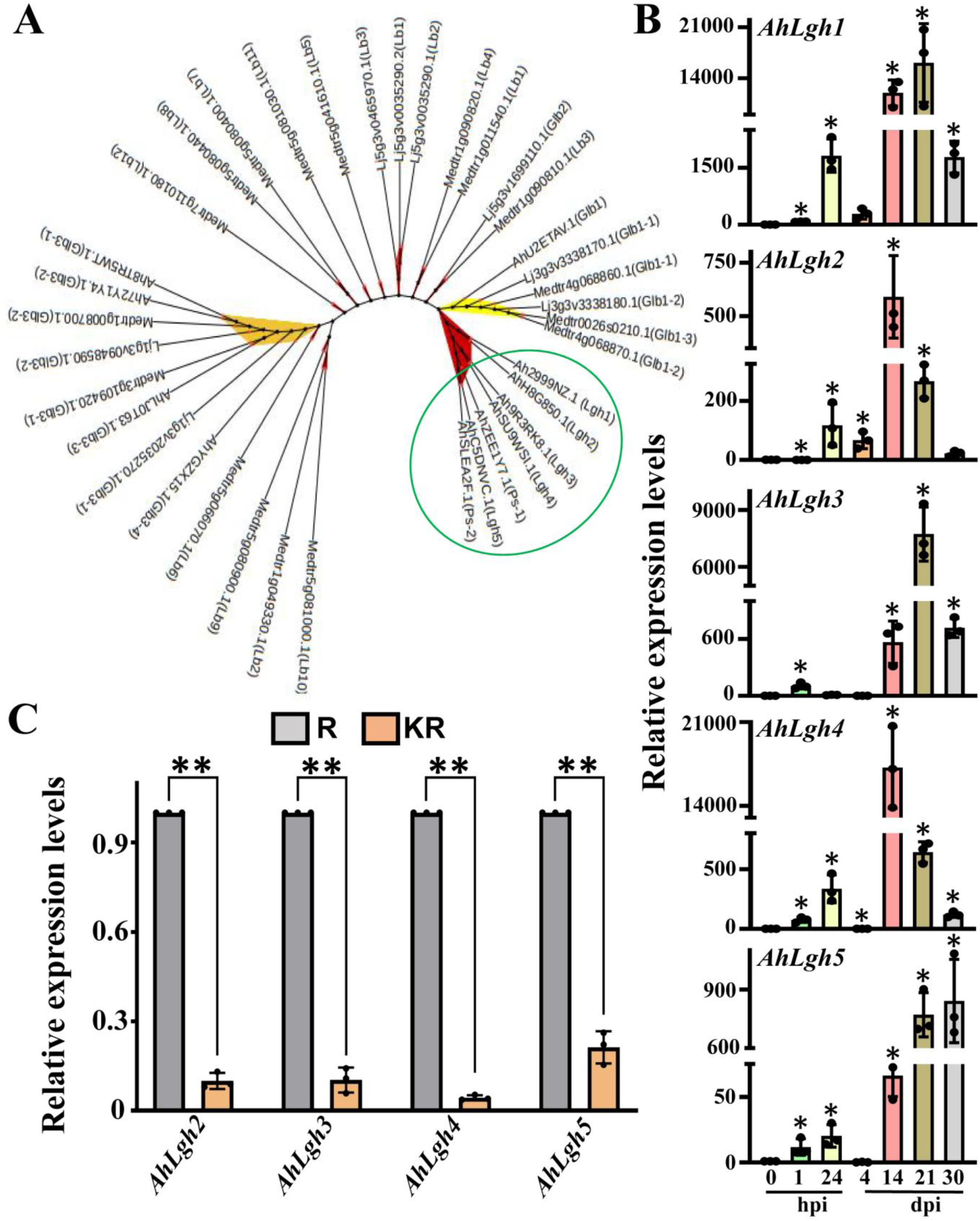
Identification and gene expression analysis of a unique group of non-symbiotic leghemoglobin genes in peanut. **A)** Phylogenetic tree of Lbs (leghemoglobin) and Glbs (globin-like protein) of *Medicago truncatula* and *Lotus japonicus* with Glbs and Lghs (non-symbiotic leghemoglobin) of *Arachis hypogaea*. A total of seven non-symbiotic leghemoglobins of *Arachis hypogaea* formed a separate group (marked within a green circle). The Glb1 and Glb3 groups were represented by yellow and orange colours, respectively. The maximum likelihood tree was constructed from a ClustalW alignment using IQ-TREE, and the final tree was visualised in iTOL. Branch support was evaluated using 1000 bootstrap replicates. **B)** Bar graphs denote the transcript abundance at different time points (hours post inoculation or hpi and days post inoculation or dpi) during nodule development compared to uninfected roots (0) by RT-qPCR. Expression in control (untreated or 0 h) was set at 1. Asterisks indicate statistically significant differences (Student’s t-test, *P < 0.01) between samples. **C)** Bar graphs denote the transcript abundance of non-symbiotic leghemoglobin genes (*AhLgh2-AhLgh5*) in nitrate plus rhizobia-treated roots (KR) compared to rhizobia-treated (R) roots at 6 dpi by RT-qPCR. The expression values were normalized against the peanut *Actin* gene expression. Expression in rhizobia-treated (R) roots was set at 1. Three biological replicates were used. Error bars represent ±SE. Asterisks indicate statistically significant differences (Student’s t-test, **P < 0.001) between samples.

Analysis of publicly available PeanutBase (https://www.peanutbase.org/) expression data set showed that all five *AhLghs* were highly expressed in nodules (Supplementary Fig. S14A), whereas except *AhGlb3-3* and *AhGlb3-4*, others *AhGlbs* were expressed in various tissues (Supplementary Fig. S14B). To verify these data, we studied the expression levels of *AhLgh* genes in four different tissues of peanut, and a significant increase in *AhLgh*s gene levels was detected in nodules compared to other tissues (Supplementary Fig. S15, A to E). We also detected a significant upregulation of *AhLgh* genes in roots at 14 dpi and 21 dpi post-rhizobial treatment (Fig. 3B). A substantial increase in the expression of all *AhLghs* at later stages after rhizobia infection suggested that they govern well-defined functions during symbiosis. Since *AhLgh1* was significantly downregulated after nitrate treatment in KR samples compared to R, we further analysed the expression of the other four *AhLghs* genes in R and KR samples to investigate whether other *AhLghs* expression was also influenced by nitrate. The expression levels of *AhLgh2*, *AhLgh3*, *AhLgh4* and *AhLgh5* were considerably affected after nitrate treatment (Fig. 3C), suggesting that nitrate transcriptionally modulates nodule-specific *AhLgh* genes during root nodule symbiosis.

### Gene silencing of *AhLgh* genes diminishes leghemoglobin content and enhances ROS production in nodules

To understand the physiological functions of *AhLgh*s in root nodule symbiosis and leghemoglobin formation, we took advantage of the hairy root transformation system using *Agrobacterium rhizogenes* and RNAi gene silencing approach. Macroscopic and microscopic observations did not disclose any noticeable alteration in root architecture. But a marked reduction (∼50%) in the number of nodule primordia formed upon inoculation with *Bradyrhizobium SEMIA 6144* at 7 dpi was detected in *AhLgh*-RNAi lines (Fig. 4A). Despite a dramatic reduction in nodule primordia number, we did not witness any major differences in the number of nodules per hairy roots at 21 dpi in *AhLgh*-RNAi lines compared to EV, as evidenced by a ∼13% less number of nodules in *AhLgh*-RNAi lines in both primary and lateral roots (Fig. 4B). According to the external view and the sectional view of nodules, the most *AhLgh*-RNAi nodules were light red in colour, whereas most EV nodules were bright red (Fig. 4C). Interestingly, we observed almost 2.5 white nodules/hairy roots in *AhLgh*-RNAi lines, which were not infected with any rhizobia and were completely absent in the EV (Fig. 4D). To evaluate the nodule colour discrepancy of *AhLgh*-RNAi and EV, we measured the amount of leghemoglobin content in root nodules spectrophotometrically. Nearly a 46% decrease in leghemoglobin content in nodules of the *Arachis hypogaea* was noticed in *AhLgh*-RNAi lines compared to EV lines (Fig. 4E). In a previous report, it was found that leghemoglobin deficiency causes a modification of the cellular redox state due to the disintegration of mitochondria (Wang et al., 2019). Thus, we studied whether silencing of *AhLgh* affected the ROS production in nodules as a result of oxidative stress. Because leghemoglobin is mainly required for detoxification of reactive oxygen species (ROS) to efficiently uphold an optimal O_2_ environment that is suitable for RNS (Hill 2012). Therefore, nodule sections were incubated with NBT and DAB to localize O_2_^•−^ free radical and H_2_O_2_ in EV and *AhLgh*-RNAi, respectively. An intense blue NBT staining was detected in *AhLgh*-RNAi nodules, whereas a very weak blue colouration was exhibited in EV nodules (Fig. 4F). In contrast to the EV nodules, which had a light reddish-brown precipitate, *AhLgh-*RNAi nodules reacted strongly with DAB to produce a robust reddish-brown precipitate (Fig. 4G). The expression of knockdown construct containing a common region of the *AhLgh*s gene produced a decrease of *AhLgh1*, *AhLgh2* and *AhLgh5* mRNA levels ranging from 90% to 95%, as compared with control empty vector (EV) roots, whereas a nearly 40% and 20% reduction of *AhLgh3* and *AhLgh4* were observed, respectively, at 21 dpi hairy roots (Fig. 4H). The phenotype exhibited by *AhLgh*-RNAi lines indicated that diminution of all leghemoglobin genes in peanut might be involved in the early phase of nodule development and leghemoglobin formation at later stages in nodules. These results confirmed that *AhLgh* genes in peanut might be involved in attenuating ROS production, which is essential for efficient nitrogen fixation.

**Figure 4.**
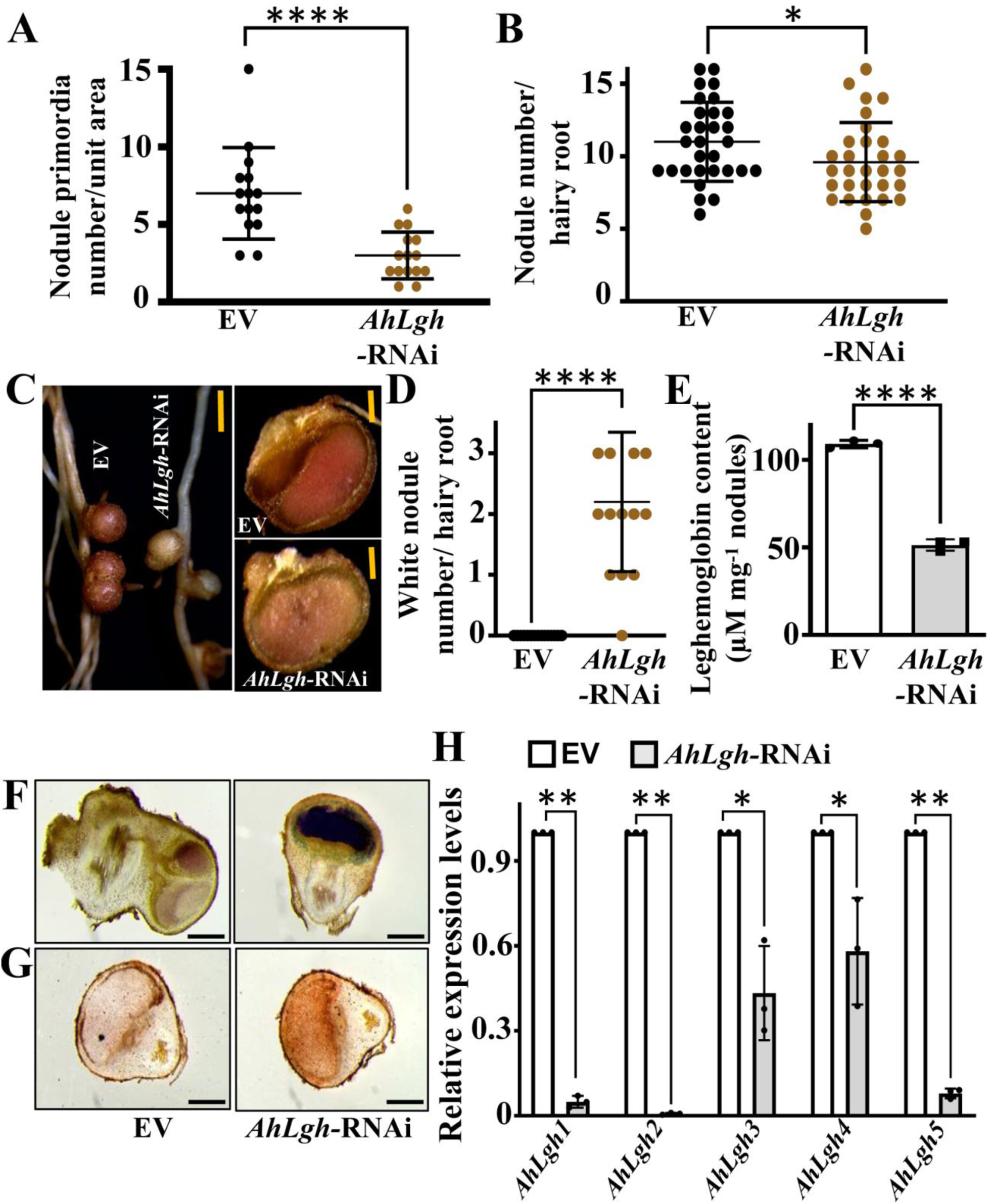
Silencing of *AhLgh* genes decreases total leghemoglobin content in nodules of peanut. **A)** Nodule primordia numbers were counted at 7 dpi in empty vector (EV) and *AhLgh*-RNAi-transformed hairy root plants inoculated with *Bradyrhizobium sp. SEMIA 6144*. **B)** Nodule numbers per hairy roots were counted at 21 dpi in empty vector (EV) and *AhLgh*-RNAi-transformed hairy root plants inoculated with *Bradyrhizobium sp. SEMIA 6144.* Three biological replicates were used, and each replicate contained 5 to 7 roots and 10 to 12 plants for the estimation of nodule primordia and nodule number, respectively. Asterisks indicate statistically significant differences (Mann-Whitney U test, ****P < 0.0001, *P < 0.05) between samples. Error bars represent ±SE. **C)** Images of intact (left panel) and transverse sections (right panel) of nodules from empty vector (EV) and *AhLgh*-RNAi-transformed hairy roots represented a difference in colour. Scale bars = 2 mm (left panel) and 0.5 mm (right panel). **D)** Quantification of white nodule numbers per hairy root in EV and *AhLgh*-RNAi transformed hairy root. Three biological replicates were used, and each replicate contained 5 to 7 roots. Asterisks indicate statistically significant differences (Mann-Whitney U test, ****P < 0.0001) between samples. Error bars represent ± standard error (SE). **E)** Estimation of leghemoglobin content in nodules of empty vector (EV) and *AhLgh*-RNAi-transformed hairy roots at 21 dpi. Asterisks indicate statistically significant differences (unpaired t test, ****P < 0.0001) between samples. Error bars represent ±SE. **F)** Transverse sections of the nodules (80 µm thickness) of empty vector (EV) and *AhLgh*-RNAi-transformed hairy roots, obtained using a vibratome, were stained with NBT (nitroblue tetrazolium) to detect the superoxide radicals. Scale bars = 0.5 mm. **G)** Transverse sections of the nodules (80 µm thickness) of empty vector (EV) and *AhLgh*-RNAi-transformed hairy roots, obtained using a vibratome, were stained with DAB (3, 3’-diaminobenzidine) to detect the endogenous hydrogen peroxide. Scale bars = 0.5 mm. **H)** Transcript levels of *AhLgh1, AhLgh2*, *AhLgh3*, *AhLgh4* and *AhLgh5* were detected by RT-qPCR in transgenic hairy roots of EV and *AhLgh–RNAi* lines at 21 dpi. The expression values were normalized against the peanut *Actin* gene expression. Expression in EV roots was set at 1. Three biological replicates were used. Error bars represent ±SE. Asterisks indicate statistically significant differences (Student’s t-test, **P < 0.001, *P < 0.05) between samples.

To further investigate the importance of non-symbiotic leghemoglobin in peanut, we generated composite plants by hairy root transformation, where overexpression of *AhLgh1* gene was driven by the CvMV promoter. Overexpression of *AhLgh1* resulted in more dark nodules and an increase in 35% leghemoglobin content in nodules (Supplementary Fig. S16, A and B). During screening of nodule phenotype, we did not observe any notable difference in nodule primordia and root nodule number at 7 and 21 dpi, respectively (Supplementary Fig. S16, A and B). Overexpression of *AhLgh1* gene was verified by RT-qPCR, and we found a significant upregulation (∼ 60-fold) of *AhLgh1* transcript in 21 dpi nodule samples compared to that of EV. These results suggested that a special group of non-symbiotic leghemoglobin genes regulate leghemoglobin content in peanut.

### NIN is required for the enhancement of leghemoglobin expression along with its canonical function

Several studies on nodule-inception (NIN) loss-of-function alleles and the cis-regulatory elements in the promoter of NIN revealed that NODULE INCEPTION (NIN), a nodulation-specific transcriptional regulator, is specifically involved in the initiation of rhizobial infection and nodule organogenesis (Schauser et al., 1999; Marsh et al., 2007; Singh et al., 2014). According to our transcriptome data, rhizobia-mediated enhancement and nitrate-dependent suppression of two *NIN* genes (*Arahy.I65W25, AhNIN1* and *Arahy.V4BGUX, AhNIN2*) overlapped with the expression of *AhLgh1* (*Arahy.2999NZ.1*). This context raised a query about whether NINs modulate leghemoglobin content in peanut. Our BLAST searches in the recently annotated peanut database (https://www.peanutbase.org/) also identified these two *NIN* genes (*Arahy.I65W25* and *Arahy.V4BGUX*). To pinpoint the impact of NIN on the expression of leghemoglobin in peanut, we employed a gene silencing approach by selecting a coding region of the gene which was similar in both the *NIN* genes. A striking difference in nodule primordia at 7 dpi and nodule number at 21 dpi in *AhNIN*-RNAi compared with the EV. Nearly 62% of the nodule primordia number and ∼75% of the nodules diminished in *AhNIN*-RNAi lines at 7 dpi and 21 dpi, respectively (Fig. 5, A to C). Similar to *AhLgh*-RNAi lines, the peripheral (external) and sectional views also exhibited a loss of brightness of red colour in *AhNIN*-RNAi, where almost 100% nodules were bright red in EV lines (Fig. 5C). One or two white nodules were occasionally observed in *AhNIN-RNAi* lines, which were entirely absent in EV lines (Fig. 5D). The variation of nodule colour in *AhNIN-RNAi* and EV lines was corroborated by measurement of leghemoglobin content, and nearly 45% leghemoglobin content was reduced in *AhNIN-RNAi* lines compared to EV lines (Fig. 5E). Whether the loss of leghemoglobin content in *AhNIN-RNAi* lines had any impact on ROS accumulation in nodules, like nitrate treatment and silencing of *AhLgh*s, we examined ROS levels by NBT and DAB staining to detect O_2_^•−^ free radical and H_2_O_2_, respectively. Similar to *AhLgh-*RNAi, an enormous accumulation of O_2_^•−^ free radical and H_2_O_2_ was detected in *AhNIN-RNAi* lines compared to EV lines (Fig. 5, F and G). To understand functional relevance, we quantified mRNA levels of both *AhNIN* genes in RNAi lines. Interestingly, the *AhNIN1* and *AhNIN2* gene expression levels at 21 dpi in *AhNIN-RNAi* lines were much lower (˂80%) than those in the EV roots, suggesting that all phenotype was the result of the inhibition of *AhNINs* (Fig. 5H).

**Figure 5.**
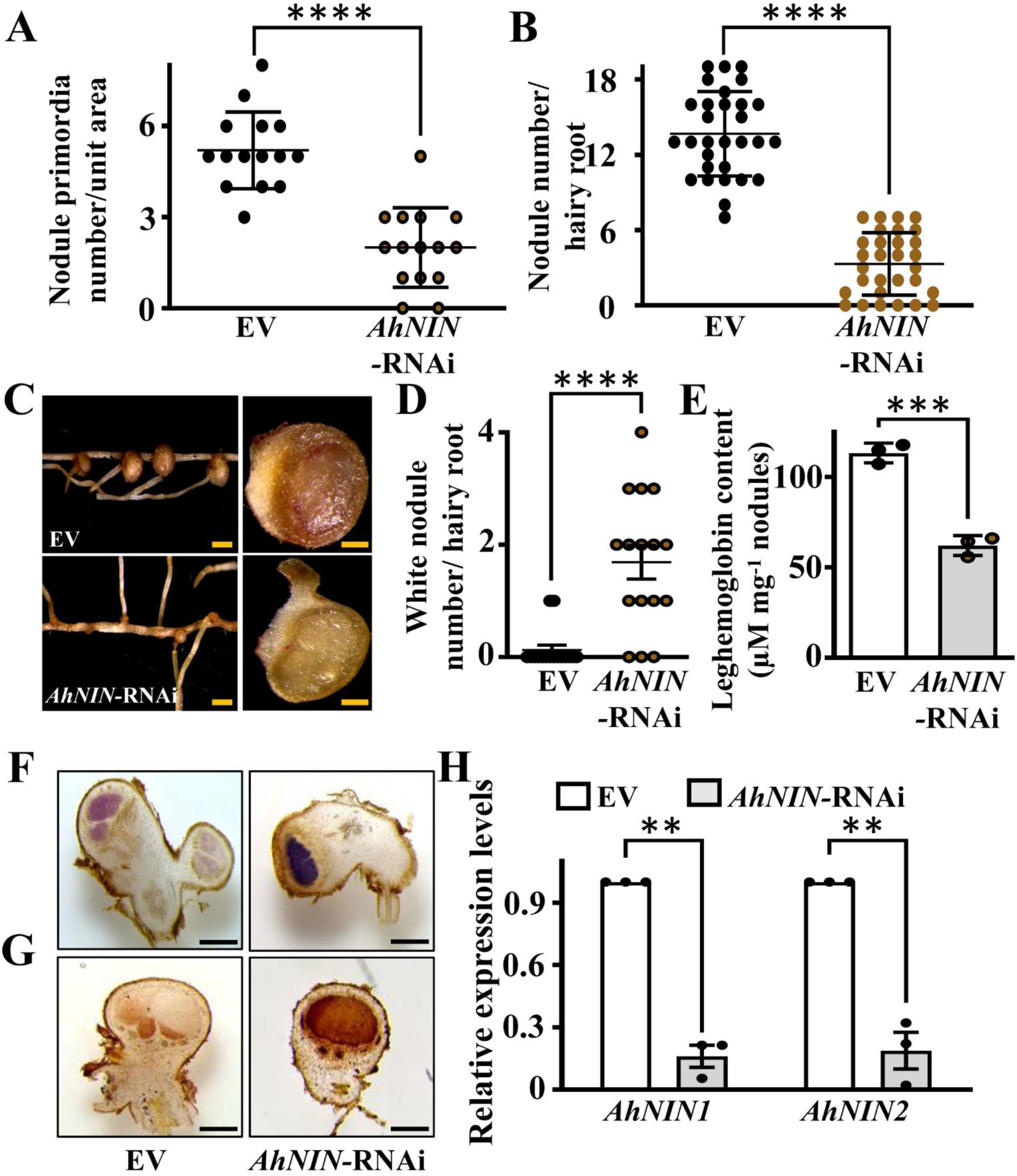
Silencing of *AhNIN* genes affected nodule organogenesis and leghemoglobin content in nodules in peanut. **A)** Nodule primordia numbers were counted at 7 dpi in empty vector (EV) and *AhNIN*-RNAi transformed hairy root plants inoculated with *Bradyrhizobium sp. SEMIA 6144*. **B)** Nodule numbers per hairy roots were counted at 21 dpi in empty vector (EV) and *AhNIN*-RNAi transformed hairy root plants inoculated with *Bradyrhizobium sp. SEMIA 6144.* Three biological replicates were used, and each replicate contained 5 to 7 roots and 10 to 12 plants for the estimation of nodule primordia and nodule number, respectively. Asterisks indicate statistically significant differences (Mann-Whitney U test, ****P < 0.0001) between samples. Error bars represent ±SE. **C)** Images of intact (left panel) and transverse sections (right panel) of nodules from empty vector (EV) and *AhNIN*-RNAi transformed hairy roots represented a difference in colour. Scale bars = 2 mm (left panel) and 0.5 mm (right panel). **D)** Quantification of white nodule numbers per hairy root in EV and *AhNIN*-RNAi transformed hairy root. Three biological replicates were used, and each replicate contained 5 to 7 roots. Asterisks indicate statistically significant differences (Mann-Whitney U test, ****P < 0.0001) between samples. Error bars represent ±SE. **E)** Estimation of leghemoglobin content in nodules of empty vector (EV) and *AhNIN*-RNAi-transformed hairy roots at 21 dpi. Asterisks indicate statistically significant differences (unpaired t test, ***P < 0.0003) between samples. Error bars represent ± standard error (SE). **F)** Transverse sections of the nodules (80 µm thickness) of empty vector (EV) and *AhNIN*-RNAi-transformed hairy roots, obtained using a vibratome, were stained with NBT (nitroblue tetrazolium) to detect the superoxide radicals. Scale bars = 0.5 mm. **G)** Transverse sections of the nodules (80 µm thickness) of empty vector (EV) and *AhNIN*-RNAi-transformed hairy roots, obtained using a vibratome, were stained with DAB (3, 3’-diaminobenzidine) to detect the endogenous hydrogen peroxide. Scale bars = 0.5 mm. **H)** Transcript levels of *AhNIN1* and *AhNIN2*, were detected by RT-qPCR in transgenic hairy roots of EV and *AhNIN–RNAi* lines at 21 dpi. The expression values were normalized against the peanut *Actin* gene expression. Expression in EV roots was set at 1. Three biological replicates were used. Error bars represent ±SE. Asterisks indicate statistically significant differences (Student’s t-test, **P < 0.001) between samples.

Similar phenotypic features, including nodule primordia number, leghemoglobin content and ROS accumulation in nodules, were observed in *AhLgh-*RNAi and *AhNIN-RNAi* lines. Therefore, it is essential to interrogate whether there is any genetic connection between *AhLghs* and *AhNINs*, which is entirely puzzling. To determine this unprecedented query, we examined the expression of five *AhLgh* genes in roots of *AhNIN-RNAi* at 21 dpi. A massive loss of *AhLghs* transcripts was detected in *AhNIN-RNAi* lines compared to EV, which was about 80% to 90% in *AhLgh1*, *AhLgh2* and *AhLgh5,* and around 40% and 60% in *AhLgh3* and *AhLgh4*, respectively (Supplementary Fig. S17A). In contrast, there was no significant difference between EV and *AhLgh-*RNAi lines in the expression of *AhNIN1* and *AhNIN2* genes examined, indicating that *AhNINs* might act upstream of *AhLgh* genes in the symbiosis signaling cascade of peanut (Supplementary Fig. S17B). Overall, it was established that *NIN* regulates nodule organogenesis and leghemoglobin synthesis in peanut during RNS.

### NIN directly upregulates non-symbiotic *Lgh* genes in peanut

To understand how NIN is involved in driving the non-symbiotic leghemoglobin expression to enhance nitrogen fixation capacity in nodules, we performed DAP-seq to pursuit the regulatory targets of *AhNIN1* at the whole-genome scale. To accomplish this, we used the genomic DNA of *Arachis hypogaea* and purified the recombinant C-terminal AhNIN1 protein, which contains a DNA-binding RWP-RK domain and a protein-protein interacting PBI domain. A total of 29,312 binding peaks were identified after overlapping two biological repeats (NIN1_1 and NIN1_2), and a total of 3,787 enriched peaks were found in the promoter, indicating strong DNA binding activities (Fig. 6, A and B). Besides 3,530; 6,712; 771; 1,485; 2,644 and 80,768 binding peaks were detected in exon, intron, 5’UTR, 3’UTR, downstream and distal intergenic region, respectively (Fig. 6B). The existence of an immense number of distal intergenic regions might be a reason for the lack of proper annotation of the peanut genome and genome duplication events in peanut. The motif prediction analysis using Find Individual Motif Occurrences (Grant et al., 2011), identified sequences related to Motif 1, which contains a 15-bp conserved sequence (5′-WHHYSAAARGKCAHW-3′) (Fig. 6C). Motif 1 was positioned mainly at the summit of the AhNIN binding peaks, thereby representing its high reliability (Figure 6D). Motif 1 was enriched with TCXCC at the 5′ side and AAAGGXCA-rich nucleotides at the 3′ side. We referred to this hallmark-containing site as NIN RESPONSIVE CIS ELEMENT (NRCE). Besides the most enriched Motif 1, it also identified Motif 2 with larger E-values (5’-TTTTACMTKTHCYWT-3’). But only Motif 1 resembled the binding motifs of NIN identified in previous ChIP-seq data of LjNIN that was previously provided using the *L. japonicus* genome version 2.5 (Soyano et al., 2014), with a minor difference. Therefore, it encouraged us to observe the interactions between AhNINs and gene promoters based on Motif 1. To better understand NIN-targeted positive regulators of root nodulation that were modulated by nitrate treatment, we compared 3,463 (after removing duplicates from 3,787 elements) found in the promoter with KR vs R downregulated genes. Our analysis identified a total of 246 genes, which were probable targets of NIN during root nodule symbiosis. We have identified *AhLgh1* (arahy.Tifrunner. gnm1.ann1.2999nz.1) as a probable target of AhNIN, which was downregulated after nitrate treatment (Fig. 6E). Interestingly, AhNIN binding site (NRCE), i.e. Motif 1, was identified in the promoter region of five *AhLgh* genes in our DAP seq analysis. The binding site was at-126 of *AhLgh1*,-35 of *AhLgh2*, – 43 of *AhLgh3*, – 89 of *AhLgh4* and –359 of *AhLgh5* from transcription start site (TSS) (Fig. 6F; Supplementary Fig. S18, A to D; Supplementary Table S9). Additionally, an intronic binding site was spotted at 2,002 of *AhLgh2*. These data suggested that *AhLgh* genes are the probable target of AhNIN. To validate *AhLgh1* is a transcriptional target of AhNIN, we performed an electrophoretic mobility shift assay using the core motif in the promoter region of *AhLgh1* as a DNA probe. Our experimental data evidently portrayed that both AhNIN1 and AhNIN2 were able to bind with the core promoter motif of *AhLgh1*, but not to the mutant probe (Fig. 6, G and H). To further corroborate *in planta*, we performed a transactivation assay in *N. benthamiana* leaves. The constructs containing *proAhLgh1*:*GUS* and empty vector (EV) or *AhNIN1* or *AhNIN2* were co-infiltrated into the leaves of *N. benthamiana*. The leaves overexpressing *AhNIN1 and AhNIN2* exhibited much stronger GUS activity than those expressing the empty vector (Fig. 6I; Supplementary Fig. S19). Altogether, these results exhibited that AhNIN1 TFs regulate leghemoglobin content by directly binding and activating the expression of *AhLgh1*.

**Figure 6.**
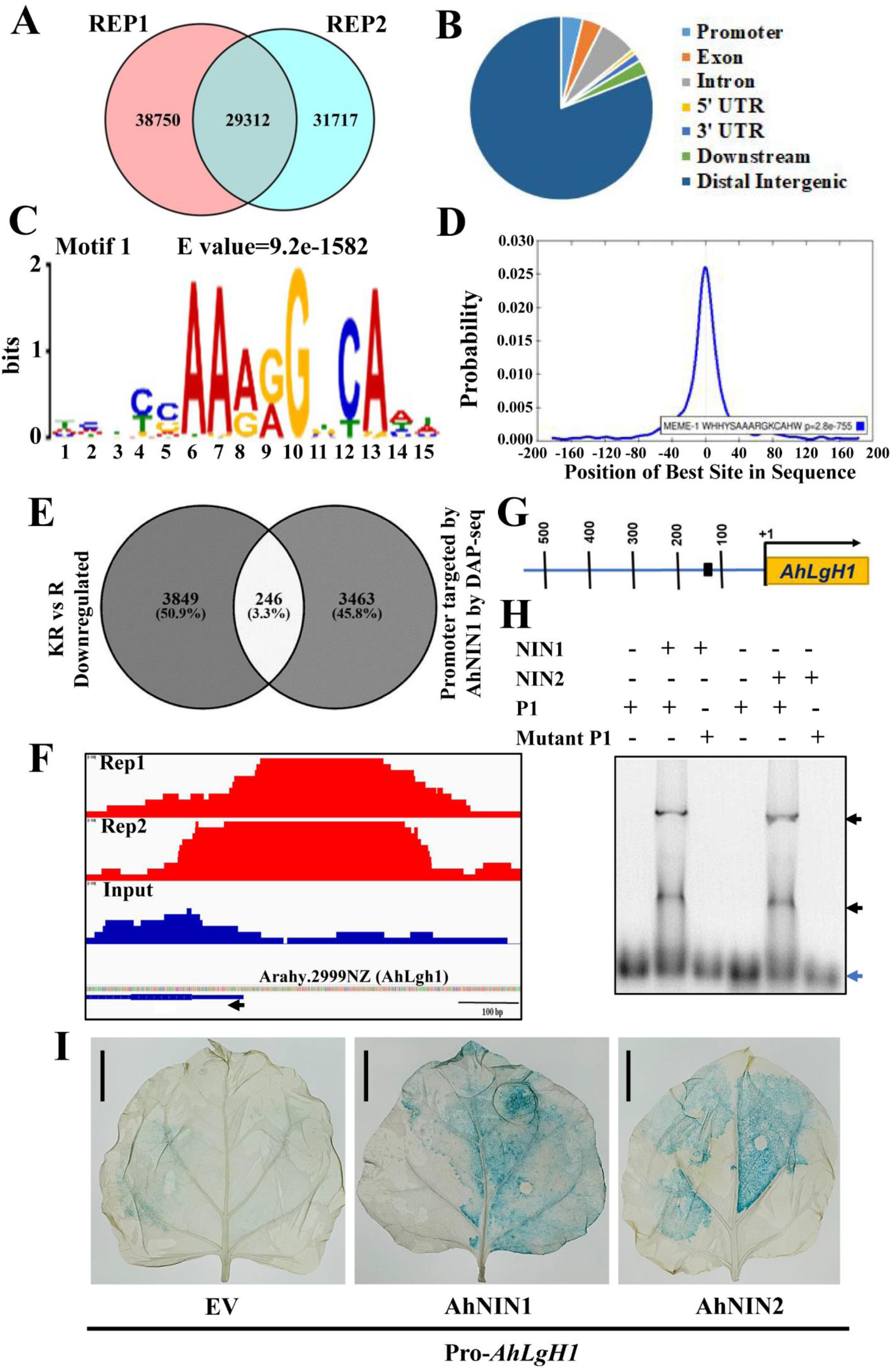
Identification of AhNIN-regulated genes by DNA affinity purification sequencing (DAP-seq). **A)** Identification of AhNIN binding sites by DAP-seq. Genomic DNA fragments (200-400 bp) from *Arachis hypogaea* roots were used for binding with *in vitro* expressed Halo-AhNIN1 C-terminal fusion protein. Comparing the sequencing results between eluates of the binding assay and input of genomic fragments, AhNIN1 binding peaks were recognized. The Venn diagram designated the number of overlapping peaks between two biological replicates (REP1 and REP2). **B)** Genome-wide distribution of AhNIN binding sites, which includes promoter, exon, intron, 5’UTR, 3’UTR, downstream and distal intergenic region. **C)** Enriched motif within the AhNIN1 binding peaks. The most significantly enriched core sequence is represented by Motif 1, consisting of 15 nucleotides. The Meme was used to calculate the *E*-value, which is 9.2e-1582. **D)** Centrimo analysis of Motif 1 displaying the positional distribution of motifs relative to the centres of DAP-seq peaks. The plot represents the average density of motif position across all motif-containing regions, spanning from −200 to +200 base pairs relative to the DAP-seq peak summit. **E)** Venn diagram of overlapping genes in two datasets: differentially expressed genes downregulated in KR vs R and promoter of genes targeted by AhNIN1 determined by DAP-seq. **F)** Localization of the enriched binding peaks of AhNIN in the promoter regions of leghemoglobin genes; *AhLgh1*. The black arrow indicates the translation start site and its orientation. Two replicates were used. Input: sequencing results of genomic DNA fragments used as a negative control. **G)** Schematic image of *AhLgh1* promoter showed the location of NIN RESPONSIVE CIS ELEMENT (NRCE). This region was determined according to the location of the binding peaks in DAP-seq results. **H)** Electrophoretic mobility shift assay (EMSA) of the interactions between AhNIN1 and AhNIN2 with the NRCE binding site of the *AhLgh1* promoter. Oligonucleotides (35 nt) were labelled and used as a probe (P1), and the mutated oligonucleotide probe (mutated P1) was used as a control. The black arrows indicate the bound probes, and the blue arrow indicates free probes. **I)** Transactivation assay in *N. benthamiana* leaves. The *GUS* (β-glucuronidase) expression driven by the *AhLgh1* promoter was enhanced by *AhNINs* (*AhNIN1* and *AhNIN2*). At least two independent leaves were examined, and three biological replicates were used. Bar=1 cm.

## Discussion

### Nitrate in soil negatively modulates root nodule symbiosis in peanut

Nitrate, a key form of inorganic nitrogen, has a multifaceted effect on plant growth and development (Vidal et al., 2014; Fredes et al., 2019; Vega et al., 2019). Nitrate availability in soil is one of the vital environmental cues that negatively regulate nodulation (Streeter 1985; Van Noorden et al., 2016; Lin et al., 2021). The high nitrate levels stimulate lateral root growth, whereas low nitrate levels preferentially allow nodule formation in soybean (Saito et al., 2014). But in peanut, high nitrate levels specifically enhanced primary root growth without altering lateral root growth. Although molecular genetic studies in model legumes established a complex nitrogen-dependent regulatory network regulating RNS in legumes to balance nitrogen requirement and carbon allocation, the intricate molecular mechanism by which nitrate diminishes efficient legume-rhizobia symbiosis is not fully understood in legumes, specifically the crack entry model peanut. According to our results, 20 mM nitrate treatment significantly decreased nodule primordia and nodule number in peanut (Fig. 1, A to C), and no nodules were formed after treatment with 40 mM nitrate (data not shown). Apart from inhibition of nodule formation, rhizobial loading was attenuated in nitrate-treated nodules. The colour changes of nodules are mainly caused by the demolish of leghemoglobin (Dupont et al., 2012). The nitrate accelerated the degradation of leghemoglobin in nodules of *Lotus japonicus* (Zhou et al., 2023). Notably, most nitrate-treated peanut nodules were less red compared to wild-type nodules, supported by ∼ 40% loss of leghemoglobin content in response to 20 mm nitrate (Fig. 1F). This might be a reason for decreased oxygen permeability and a reduced supply of oxygen to bacteroids, which eventually inhibits nitrogenase activity prerequisite for nitrogen fixation (Becana and Sprent Becana 1987). Nitrate also causes an inhibition of ROS content in the primary root tip (Zang et al., 2020), but unexpectedly, we found an enormous accumulation of O_2_^•−^ free radical and H_2_O_2_ in nitrate-treated peanut nodules, suggesting that an enhanced accumulation of reactive oxygen species occurred in response to nitrate (Fig. 1, G and H). Furthermore, the high nitrate content in KR roots strengthened these phenotypic consequences, resulting from the accumulation of nitrate. Various phenotypic alterations of nodules upon nitrate treatment prompted us to perform transcriptome analysis to understand the underlying mechanisms of nitrate effects on nodule formation in peanut.

Our study aimed at identifying gene regulatory cascades to recognize the impact of nitrate on root nodulation in roots during the early stages of symbiosis. The significant enrichment of mitochondria-related cellular components and ATP-binding related molecular functions in upregulated R and downregulated KR indicated that the potential modulation of leghemoglobin, which is a prerequisite for providing oxygen to mitochondria to maintain cellular respiration and nitrogen incorporation in the symbiosome. According to KEGG pathway, the nitrate-mediated diminution of rhizobia induced genes related to hormone signaling, MAPK signaling, plant pathogen interaction, nitrogen metabolism etc. were similar to the mechanism related to cessation of nodulation in any legumes (Fig. 2B). Based on the complex effect of nitrate treatment at the early stages of nodulation by modulating the expression of several genes, numerous metabolic pathways and multiple transcription factors, a full uncoupling of nitrate availability and nitrogenase activity was possible in peanut. The impact of nitrate was demonstrated in the transcriptome of model legumes, like *Lotus japonicus* and *Medicago truncatula,* by abrogation of several symbiotic genes related to senescence, ammonium transporter or channel in the symbiosome membrane, nodulins and respiratory genes of nodules (Cabeza et al., 2014; Wang et al., 2023). By combining the genes that are upregulated after rhizobia treatment (R) and genes that are downregulated in rhizobia plus nitrate treatment (KR), we have identified 10 symbiotic genes in peanut that were closely associated with nodulation signaling cascades. Out of 10, two were *AhNIN* genes and one was leghemoglobin (*AhLgh1*), indicating these rhizobia inducible genes were inhibited in response to nitrate (Fig. 2, C and D). Nitrate-based transcriptional inhibition of *AhNINs* in peanut is fascinating, as only the functional activity of NIN is compromised in NLP-dependent manner in model plants (Lin et al., 2018b).

### Non-symbiotic leghemoglobin, an essential component of root nodule symbiosis in peanut, is inhibited by nitrate treatment

Leghemoglobin, one of the predominant heme-containing oxygen-carrying proteins of mature nodules in legumes (Marcker et al., 1984; Ott et al., 2005). To make efficient nitrogen fixation, leghemoglobin maintains an optimal O_2_ environment inside cells of nodules and delivers it to mitochondria for cellular respiration. In return, rhizobia synthesize heme for proper nitrogen fixation of legumes (O’Brian 1996; Smagghe et al., 2009; Vázquez-Limón et al., 2012). Similar to the other legumes that followed the infection thread model, the crack entry model peanut genome is encoded by multiple leghemoglobins (AhLghs) (Ott et al., 2005; Smagghe et al., 2009; Larrainzar et al., 2020). Evolutionary analysis revealed that five leghemoglobin (AhLghs) proteins exhibit a distinct group of nodule-specific non-symbiotic leghemoglobin, which were different from globin-like proteins (one AhGlb1 and four AhGlb3) based on protein motif architecture and *in silico* structural alignment (Fig. 3A; Supplementary Fig. S11 and S12). The intense sequence similarity and resemblance of gene and motif architecture among the five leghemoglobin (AhLghs) demonstrated their role as the leghemoglobin in peanut. Transcript analysis revealed that all five leghemoglobins were mainly nodule-specific proteins, and their expression was induced in the root at later stages of rhizobial treatment in peanut (Fig. 3B). In *Medicago sativa*, transcripts of leghemoglobin genes were first detected 9-10 days after rhizobium inoculation, and subsequently, increased and remained at steady-state levels during nodule development (Barker et al., 1988). The promoter of the leghemoglobin gene of *Vicia faba* (*VfLb29*) was precisely active in the nitrogen-fixing zone of root nodules and arbuscule-containing cells after rhizobial and endomycorrhizal fungus infection, respectively (Vieweg et al., 2004). Gene silencing of five leghemoglobins in peanut exhibited a drastic reduction of leghemoglobin content in nodules without affecting nodule number and root phenotype under symbiotic conditions, underpinning the importance of *AhLghs* for nitrogen fixation during root nodule symbiosis but not for plant developmental cues. Although at initial developmental stages, nodule primordia numbers were less in *AhLgh* RNAi lines compared to EV lines, later these RNAi lines showed almost equal numbers of nodules, like EV (Fig. 4, A to E). In Lotus, RNAi suppression of three leghemoglobin genes severely affected symbiotic nitrogen fixation by producing small white nodules under symbiotic conditions (Ott et al., 2005). We also detected an increased number of white nodules in peanut *AhLgh* RNAi lines, but the RNAi phenotype was not as profound as that of lotus under symbiotic conditions, presumably due to an inadequate inhibition of *AhLgh3* and *AhLgh4* or the functional divergence of leghemoglobin of the crack entry model peanut from the infection thread model following legumes.

The treatment of nitrate had a significant effect on the expression of *AhLgh* genes, like other nodulation-specific genes, based on transcriptome analysis. Interestingly, a significant suppression of *AhLgh1*, *AhLgh2* and *AhLgh5* upon nitrate treatment indicated that *AhLgh1*, *AhLgh2* and *AhLgh5* were primary targets of nitrate in peanut. In contrast, a partial suppression of *AhLgh3* and *AhLgh4* was observed upon nitrate treatment (Fig. 2E and 3C). The transcript pattern of *AhLghs* was in line with the changes of leghemoglobin content in response to nitrate. In addition, the expression pattern of the five *AhLgh* genes upon nitrate treatment was somewhat similar to *AhLgh* RNAi lines, which might be a plausible indication of comparable phenotypic outcomes. Our investigation also suggested that nitrate treatment endorsed a genetic machinery to suppress the expression of five *AhLgh* genes, thereby decreasing the amount of leghemoglobins, essential for symbiotic nitrogen fixation. Similar to nitrate treatment, the accumulation of ROS, like O_2_^•−^ free radical and H_2_O_2,_ was detected in mature nodules of *AhLgh* RNAi lines, signifying that inhibition of leghemoglobin genes altered the cellular redox state in nodules probably as a result of oxidation of redox-sensitive proteins like nitrogenase or ferredoxin (Fig. 4, F and G).

### Nitrate hinders NIN-dependent transcriptional regulation of leghemoglobin to fix nitrogen

NIN TF is derived from the duplication event of NLP by amino acid substitutions, and this event highlights their adaptive significance to drive the nitrogen-fixing nodule (NFN) symbiosis in legumes during evolution (Zhang et al., 2024). The predominant functional role of NIN related to rhizobial entry, nodule organogenesis, nitrogen fixation and auto-regulation of nodulation is well established in model legumes that follow the infection thread model (Shen and Feng 2023). But in peanut, a crack entry model, NIN is not enabling rhizobial entry as its expression remains only in the root pericycle and cortex (Bhattacharjee et al., 2022). To understand how NIN modulates RNS in peanut, we identified two NIN genes (*AhNIN1* and *AhNIN2*) from the peanut by searching the recently annotated genome, and both nodule primordia and nodule formation were strikingly inhibited by silencing of *NIN* genes, indicating their utmost role in nodule organogenesis in peanut (Fig. 5, A and B). The loss of leghemoglobin content and accumulation of ROS in nodules of gene-silencing lines of *AhNIN* suggested their pivotal role in leghemoglobin formation, thereby contributing to nitrogen fixation in RNS (Fig. 5, E to G). These data demonstrate that both NINs from peanut have conserved functions for driving nodule development and nitrogen fixation. To explore in detail the NIN-mediated transcriptional regulation in the crack entry model peanut, we identified multiple genes as candidates for AhNIN direct targets by DAP-seq analysis. Comparison of our transcriptome and DAP seq data predicted that rhizobia-induced 246 genes were transcriptionally inhibited by nitrate (Fig. 6, A to E). Interestingly, we found all five leghemoglobin genes within that gene pool, which was supported by the decline of leghemoglobin content in the nodule under nitrate treatment (Fig. 1F).

The transcriptional regulation of leghaemoglobin is a sophisticated and tightly regulated process. In the model legume *Medicago truncatula*, NIN-like (NLPs) TFs regulate leghemoglobin genes to enhance nitrogen fixation (Jiang et al., 2021). According to our DAP seq data, in the peanut that followed the crack entry model, leghemoglobin genes were direct targets of NIN, as all five leghaemoglobin promoters contained NRCE or motif 1. We corroborated this data by EMSA and transactivation assay using AhLgh1 promoter (Fig. 6, H and I). Further downregulation of five leghaemoglobin genes in *AhNIN*-RNAi lines, implying that *AhLghs* in peanut are transcriptionally regulated by AhNIN. In contrast, no changes in *AhNINs* expression were detected in *AhLgh*-RNAi lines. On the other hand, according to our transcriptome and gene expression data, nitrate treatment decreases the expression of both *AhNINs* and *AhLghs*. Leghemoglobin content was one of the vital factors that declined under nitrate treatment and both gene silencing lines of *AhNINs* and *AhLghs*. Concurrently, an enhanced ROS accumulation was observed due to misaccumulation of leghemoglobin under nitrate treatment in both gene silencing lines of *AhNINs* and *AhLghs*.

Based on our findings, we propose a model illustrating that nitrate-dependent inhibition of NIN reduces leghemoglobin, thereby impeding the scavenging of oxygen or oxygen buffering, essential for nitrogen fixation in peanut. Based on previous studies, we can predict a plausible scenario where nitrate-induced NLPs can hinder the binding of AhNIN to the AhLgh promoter, because the consensus of the NRCE sequence is very similar to the NLP binding site. In *Medicago truncatula*, nitrate signal promotes accumulation of NLP1 in the nucleus, probably via nitrate-CPK-NLP signaling cascades, for transcriptional inhibition of NIN via direct binding or competing for binding sites on the promoters of the NIN target genes (Liu et al., 2017; Lin et al., 2018b). The nitrate-induced NLP4 forms a heterodimer with NIN to restrict the binding of NIN homodimer to its targets during transcriptional activation downstream of CLE-RS2 and NF-YB genes, thereby modulating their expression in *Lotus japonicus* (Nishida et al., 2021). Ultimately, our results underscored the fluctuations of gene regulatory cascades under nitrate treatment. Further functional and molecular analysis of *AhNINs* and *AhLgh* revealed one important strategy where NIN can modulate *AhLgh* expression in a nitrate-dependent manner to control excess nitrogen fixation for balancing nitrogen acquisition (Fig. 7). Comprehending these kinds of mechanisms of transcriptional regulation is a prerequisite to optimising the direct pathway for nitrate uptake, contributing to advancements in nitrogen fixation.

**Figure 7.**
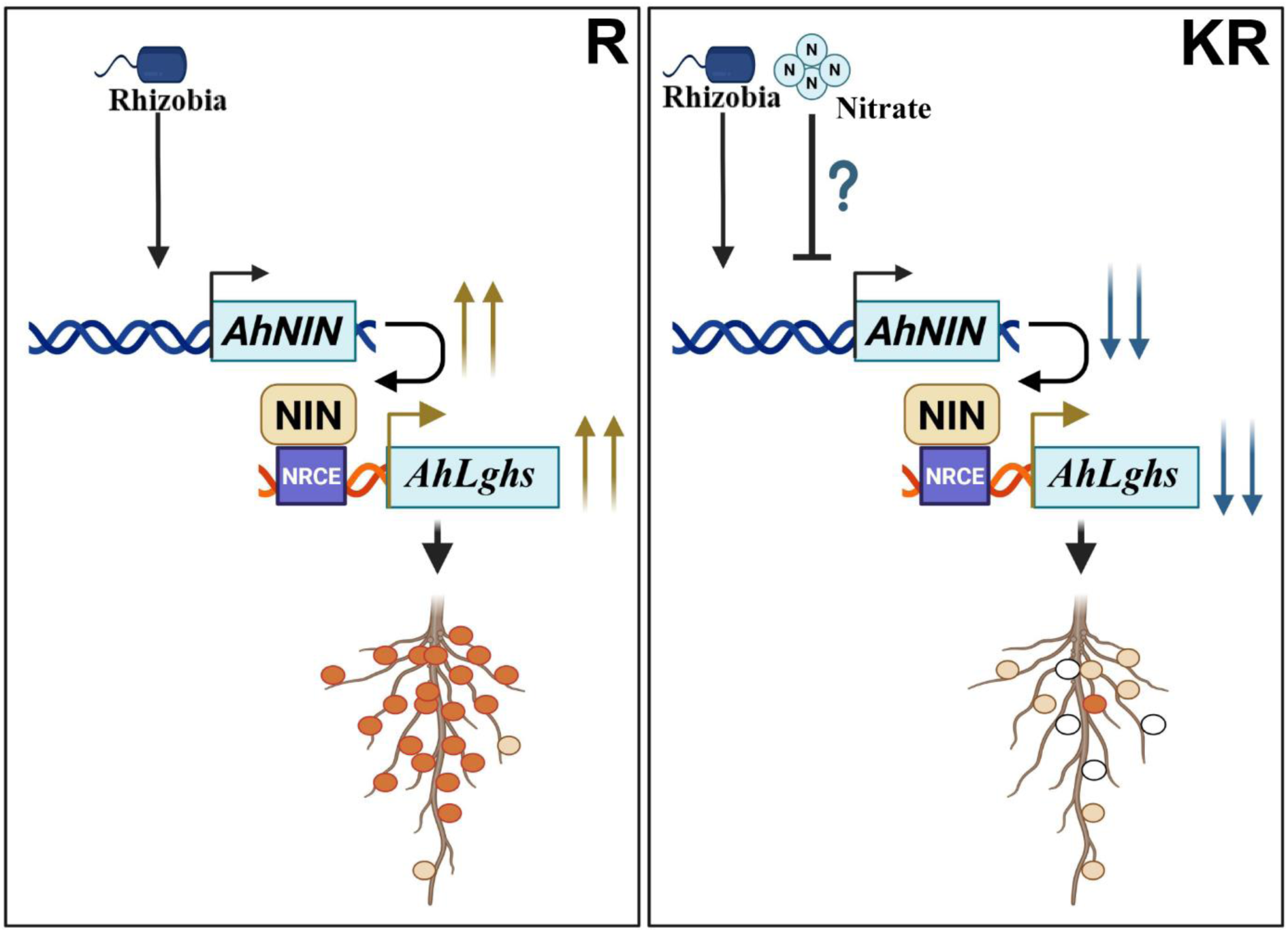
A proposed model in which nitrate-mediated downregulation of NIN affects non-symbiotic leghemoglobin genes in peanut. In the presence of rhizobia (R), the rhizobial signal induces the expression of the *NIN* gene, which encodes a transcription factor. Subsequently, NIN binds to the NIN RESPONSIVE CIS ELEMENT (NRCE) of the promoter of non-symbiotic leghemoglobin genes (*AhLghs*) to promote their expression, thereby leading to high leghemoglobin accumulation and the development of strong red-coloured nodules. By contrast, in the presence of rhizobia together with nitrate (KR), NIN expression is significantly reduced. This downregulation of NIN further decreases *AhLghs* expression, resulting in diminished leghemoglobin content within the nodules. It leads to the formation of pale red to greenish nodules, with some white nodules due to insufficient leghemoglobin accumulation.

## Materials and methods

### Plant material and bacterial strains

The seeds of peanut (*Arachis hypogaea* cultivar Kadiri-6/K-6) were collected from Acharya N.G. Ranga Agricultural University, Regional Agricultural Research Station (RARS) in Tirupati for our experiments. For hairy root transformation, *Agrobacterium rhizogenes* strain R1000 was employed (Sinharoy et al., 2009), and *Agrobacterium tumefaciens* GV3101 strain was used for transient expression analysis in *Nicotiana benthamiana*. The rhizobial strain *Bradyrhizobium* sp. SEMIA 6144 was used for root nodulation (Sinharoy et al., 2009). Primers used in this study are in Supplementary Table S10.

### Nitrate treatment

The peanut seeds were surface-sterilized by 70% ethanol for ∼2 minutes with gentle agitation, followed by rinsing with sterile deionized water. Surface-sterilized peanut seeds were soaked overnight and transferred to a pot containing sterile vermiculite, perlite and soilrite (2:1:1) under a 16-h light/8-h dark cycle, 65% relative humidity at 28°C in a growth chamber. Ten-day-old seedlings were treated with 10 ml of 10 mM and 20 mM KNO_3_ from 11 to 17 days and 25 to 31 days. Plants were inoculated with rhizobia at 21 days (Raul et al., 2022). Nodule primordia and nodule numbers were counted at 7 dpi (days post-infection) and 21 dpi. Ten ml of Nitrogen-free B&D media was added to each pot on each alternate day (Broughton and Dilworth 1971). Autoclave water was used as a mock or control. The root samples were harvested at 6 dpi from control, rhizobia-treated (R) and 20 mM nitrate plus rhizobia (KR) treated samples for RNA sequencing.

### RNA Sequencing

The total RNA was extracted from control, rhizobia-treated, and rhizobia plus nitrate-treated roots using the RNeasy Plant Mini Kit. After mRNA purification, library preparation and sequencing were carried out using the Illumina NovaSeq 6000 platform with paired-end reads (Bionivid Technology, Bengaluru). To ensure high-quality data for downstream analysis, quality control and pre-processing of FASTQ files were performed. The quality filtering and adapter trimming of raw reads were performed using Fastp (Chen et al. 2018), with a quality Phred score cutoff of 30. The Hisat2 splice-aware aligner was used to map the RNA-seq reads to *Arachis hypogaea* genome (arahy.Tifrunner.gnm1.KYV3-Genome-Assembly) (Kim et al. 2019). Transcriptome assembly and quantification for each sample were performed using StringTie (Pertea et al. 2015), and gene expression levels were normalized using FPKM (Fragments Per Kilobase of exon per Million mapped fragments) method. Functional annotation of assembled transcripts was conducted using NCBI RefSeq resources with Gene Ontology (GO) terms. Differential gene expression analysis was performed using the limma-voom package within the Galaxy platform. OmicsBox was used for the KEGG pathway analysis. (see details in Supplementary materials).

### Gene cloning, construct preparation and gene expression analysis

Total RNA was extracted using Trizol reagent (Takara), and after cDNA synthesis, diluted cDNA was used for gene amplification using gene-specific primers (Palaka et al., 2021). The full-length cDNA of *AhNIN1*, *AhNIN2* and *AhLgh1* was amplified from cDNA to clone into pCR™8/GW/TOPO™ TA cloning vector (Invitrogen) for sequence verification. The full-length *AhLgh1* gene was transferred into pCAMGFP-CvMV-GWi (CvMV promoter) binary vector from pCR8/GW/TOPO TA vector (Invitrogen) using LR Clonase (Invitrogen) for preparation of overexpression construct (Choudhury and Pandey 2013). To prepare RNAi constructs for *AhNINs* and *AhLghs*, 400 bp and 453 bp fragments were amplified from cDNA, respectively. These gene fragments were cloned into pCR™8/GW/TOPO™ TA cloning vector for sequence verification (Invitrogen) and subsequently transferred into CGT11017A binary vector by Gateway-based cloning using LR Clonase (Invitrogen). All constructs were introduced into *Agrobacterium rhizogenes* strain R1000 to generate transgenic hairy roots.

For gene expression analysis, real-time quantitative reverse transcription PCR was performed using SYBR Green (Takara). The *AhActin* gene was used as an internal control, and the 2^ΔΔCt method was used for the calculation of relative expression (Schmittgen and Livak 2001). Three biological replicates were used.

### Hairy root transformation

Peanut seeds (*Arachis hypogaea* cultivar Kadiri-6/K-6) were grown in a greenhouse (14 h light/10 h dark cycle) for 10 days at 40-60% relative humidity and 28°C. The hairy root transformation was carried out by modification of the protocol described by (Nanjareddy et al., 2022). After two weeks, plants containing transgenic hairy roots were transferred to nitrogen free autoclave soil mixture consisting of vermiculite, perlite and soilrite in a 2:1:1 ratio supplemented with Nitrogen-free B&D media under a 14-h light/10-h dark cycle, 40-60% relative humidity at 28°C in a growth chamber. Subsequently, plantlets were treated with actively growing *Bradyrhizobium sp.* SEMIA 6144 (0.1 at OD600). The number of nodule primordia and nodules was quantified at 7-and 21-day post-infection (dpi). Three biological replicates were used, and at least 10-12 transgenic hairy roots were used in each experiment.

### Microscopy

Root and nodule images were captured using an SZX7 stereomicroscope (Olympus) equipped with an XCAM 8.0MP camera and processed with the ImageView software. The roots and nodules were embedded in 2% agarose and subsequently sectioned using a vibratome (VT 1000S; Leica). Thin nodule sections were stained with toluidine blue and observed by light microscopy to differentiate rhizobia loading. The SYTO 13, a cell-permeant fluorescent nucleic acid-binding dye (Thermo Fisher Scientific), was used to visualize the development and distribution of rhizobia in root nodules by a Leica TCS SP8 spectral confocal laser scanning microscope. In nodule sections, superoxide radical and H_2_O_2_ production were determined by using nitroblue tetrazolium (NBT) and 3,3’-diaminobenzidine (DAB) staining, respectively, according to (Kaur et al., 2016; Minguillón et al., 2024).

### Leghemoglobin concentration determination

To measure non-symbiotic hemoglobin, 100 mg of fresh nodules were ground and homogenized in 0.3 ml of pre-cooled PBS buffer (Na₂HPO₄-NaH₂PO₄, pH 6.8). The homogenate was centrifuged at 12,000g for 15 min, and subsequently, the supernatant was analysed by spectrophotometry at 540, 520 and 560 nm (LaRue and Child, 1979; Jiang et al., 2021). The non-symbiotic hemoglobin content was calculated using bovine Hb as the protein standard.

### DAP Seq analysis

The extracted genomic DNA (gDNA) from (*Arachis hypogaea*) was fragmented using a Covaris M220 (Woburn, MA, USA), following the manufacturer’s instructions. After purification by Agencourt AMPure XP Kit (Beckman), adapter was ligated with fragmented gDNA. The c-terminal coding sequence of *AhNIN1* including RWP-RK and PB1 domains was cloned into pFN19K HaloTag T7 SP6 Flexi expression vector, and after protein expression using the TNT SP6 Coupled Wheat Germ Extract System (Promega), proteins were captured using Magne HaloTag Beads (Promega). The adapter-ligated gDNA fragments were incubated with protein-bound beads on a rotator for 1 h. After removing unbound DNA, bound DNA fragments were eluted for PCR with KAPA HiFi HotStart Ready-mix PCR Kit (Roche) using Illumina TruSeq Universal primer and an Illumina TruSeq Index primer. After purification, the eluted DNA fragments were sequenced on an Illumina NovaSeq 6000 platform (CD Genomics, USA). The DAP-seq libraries without adding protein-bound beads were used as negative control.

### Recombinant protein purification and electrophoretic mobility shift assay (EMSA)

The c-terminal coding sequences of *AhNIN1* and *AhNIN2* including RWP-RK and PB1 domains were cloned into pET8a expression vectors. The recombinant proteins were produced in *E. coli* strain Rosetta (DE3) (Novagen) by inducing for 12 h at 16°C using 0.5 mM isopropyl β-D-1-thiogalactopyranoside (IPTG) and subsequently purified using Ni-NTA agarose (Qiagen) according to (Choudhury et al., 2012).The concentration of the purified recombinant proteins was assessed using the Bradford method (Kielkopf et al., 2020). To prepare oligonucleotide probes for EMSA, equivalent molar amounts of two oligos were annealed by heating to 90°C for 5 minutes and subsequently, the oligos were allowed to slowly cool to room temperature (RT) over 45 minutes. The reactions containing ∼4.5 µg *AhNIN1* and *AhNIN2* C-terminal recombinant proteins, (25ng/µl) double stranded probe and binding buffer were incubated for 45 minutes at RT (25 mM HEPES-KOH, pH 8.0, 50 mM KCl, 1 mM DTT, 0.05% [v/v] Triton X-100, and 5% [v/v] glycerol) before resolving by electrophoresis in 0.5× TBE buffer. Electrophoretic Mobility-Shift Assay (EMSA) Kit, with SYBR™ Green (Thermofisher) was used for staining of gel after electrophoresis.

### Transactivation assay

To generate *proAhLgh1::GUS*, the 500 bp promoter region of *AhLgh1* was amplified and cloned into pCAMBIA1305.2 vector at upstream of the *GUS* gene before transforming into *A. tumefaciens* strain GV3101. The full-length *AhNIN1* and *AhNIN2* genes were transferred into pEarlyGate203 (CaMV 35S promoter) binary vector from pCR8/GW/TOPO TA vector (Invitrogen) using LR Clonase (Invitrogen). Both CaMV 35S:*AhNIN1/AhNIN2* and the EV (control) were individually transformed into *A. tumefaciens* stain GV3101. The Agrobacterium cells were resuspended in infiltration buffer (10 mM MgCl_2_,10 mM MES, pH 5.6, and 150 μM acetosyringone) after collection by centrifugation. Subsequently, in various combinations, Agrobacterium cells were infiltrated into *N. benthamiana* leaves, and the activity of the enzyme β-glucuronidase (GUS) was detected by a histochemical assay after 48 h according to Chen et al. (2013).

## Statistical analysis

The data presented in all figures were obtained from at least three biological replicates. To calculate statistical significance, Dunn’s multiple-comparisons test, Mann-Whitney U test and Student’s t-test were applied. All analyses were conducted using GraphPad Prism software version 8.0.2.

## Data availability

The sequence raw files from RNA sequencing were submitted to the NCBI SRA database under BioProject number PRJNA1329962 (BioSample Accessions: SRR35628999 to SRR35629010).

## Acknowledgments

This study was supported by the plant growth rooms, greenhouses and microscopy facilities of the Indian Institute of Science Education and Research (IISER), Tirupati. The authors are thankful to the members of the Plant Signalling Lab at IISER Tirupati for their support. RK acknowledges the University Grants Commission (UGC), Government of India, for Ph.D. fellowship.

## Author contributions

RK and SRC: designed the research. RK: performed the experiments. RK: analyzed the data. RK: prepared the original draft. SRC: writing, reviewing, and editing.

## Funding

The authors are grateful for funding by the ANRF Research Grant (CRG/2023/004559) of SRC and the STARS Research Grant of SRC (MoE/STARS-1/508).

## Conflict of interest statement

The authors declare no competing interests.

## Supplementary data

**Supplementary Figure S1.** Flow chart of nitrate treatment in peanut.

**Supplementary Figure S2.** Effect of different concentrations of nitrate on root architecture in peanut.

**Supplementary Figure S3.** Nitrate treatment restricts rhizobial colonization in mature nodules in peanut.

**Supplementary Figure S4.** Phenotype and biochemical features of nitrate-treated nodules in peanut.

**Supplementary Figure S5.** Transcriptomic analysis of nitrate-treated roots.

**Supplementary Figure S6.** GO analysis of common DEGs between upregulated genes in rhizobia-treated roots compared to control (R vs control) and downregulated genes in nitrate plus rhizobia-treated roots compared to rhizobia-treated roots (KR vs R).

**Supplementary Figure S7.** Co-expression network of common DEGs between upregulated genes in rhizobia-treated roots compared to control (R vs control) and downregulated genes in nitrate plus rhizobia-treated roots compared to rhizobia-treated roots (KR vs R).

**Supplementary Figure S8.** Prediction of transcription factors (TFs) within common DEGs.

**Supplementary Figure S9.** Comparative sequence analysis of non-symbiotic leghemoglobins of peanut.

**Supplementary Figure S10.** Chromosomal localization and gene architecture of non-symbiotic leghemoglobin genes of peanut.

**Supplementary Figure S11.** Phylogenetic tree and conserved motif of leghemoglobins of peanut.

**Supplementary Figure S12.** Comparative sequence analysis of AhLgh1 with AhGlb1.1 of peanut.

**Supplementary Figure S13.** The structure of non-symbiotic leghemoglobin of peanut was compared with hemoglobins of legumes and non-legumes.

**Supplementary Figure S14.** Expression profiles of non-symbiotic leghemoglobin genes in peanut.

**Supplementary Figure S15.** Tissue-specific expression of non-symbiotic leghemoglobin genes in peanut.

**Supplementary Figure S16.** Overexpression of *AhLgh1* increases leghemoglobin content.

**Supplementary Figure S17.** NIN acts upstream of *AhLgh1*.

**Supplementary Figure S18.** NIN binds to the NRCE of *AhLgh*’s promoter region.

**Supplementary Figure S19.** Quantification of *GUS* gene expression.

**Supplementary Table S1.** A complete summary of DEGs.

**Supplementary Table S2.** Common 2031 elements in R vs Control Up and KR vs R Down.

**Supplementary Table S3.** Significant GO enrichment analysis of common 2031 genes in R vs Control Up and KR vs R down.

**Supplementary Table S4.** KEGG analysis of Common 2031 genes in R vs Control Up and KR vs R down.

**Supplementary Table S5.** List of SYM Genes from R vs Control Up and KR vs R down.

**Supplementary Table S6.** Protein-protein interaction enrichment of common proteins in R vs Control Up and KR vs R down.

**Supplementary Table S7.** Common transcription factors in R vs Control Up and KR vs R down.

**Supplementary Table S8.** Structural matchmaking of AhLgh with MtLb1 and other non-legume hemoglobin like OsHb1, AtHb1 based on alignment score and RMSD.

**Supplementary Table S9.** Enriched peaks in DAP-seq global profiling of AhNIN1 binding sites (NBS).

**Supplementary Table S10.** Primers used in this study.

**Supplementary material**

## Notes

### Competing Interest Statement

The authors have declared no competing interest.

