## Supplementary Figures for "Nitrate restricts the expression of non-symbiotic leghemoglobin through inhibition of nodule inception protein in nodules of peanut (*Arachis hypogaea*)"

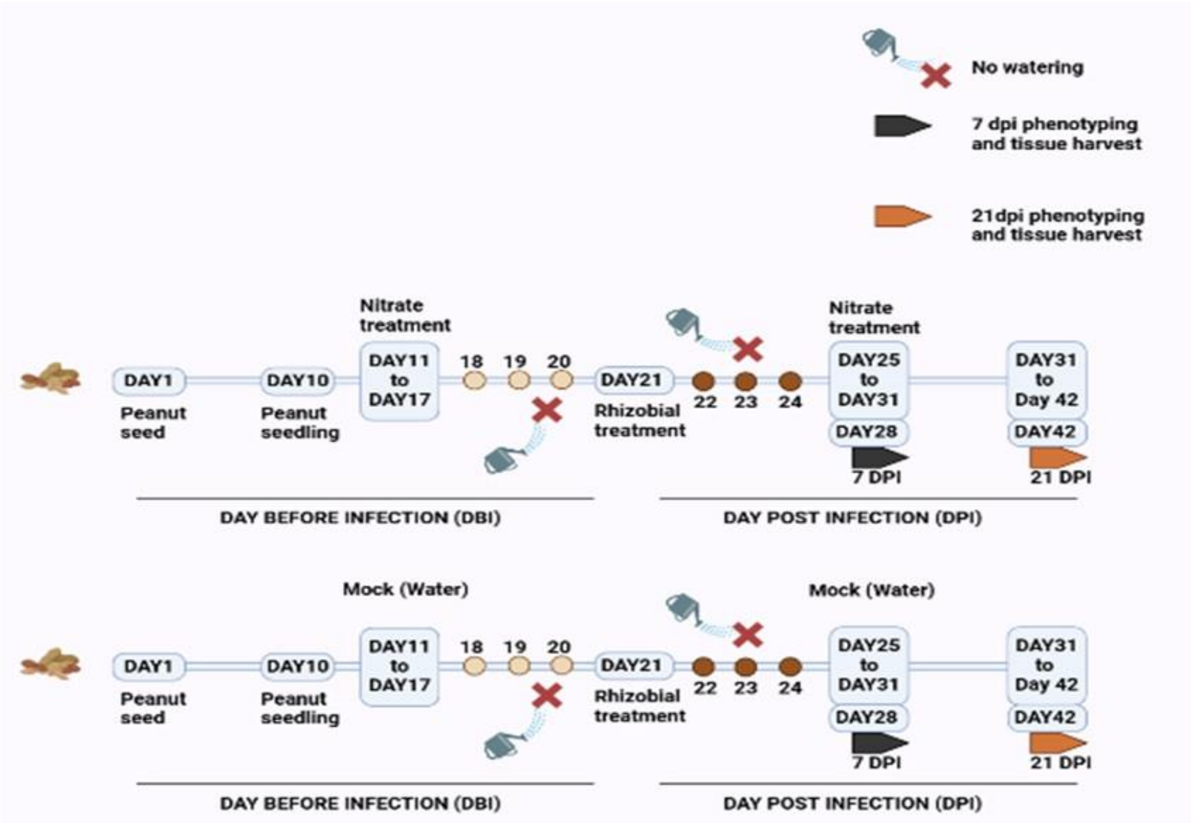


**Figure S1. Flow chart of nitrate treatment in peanut.** Peanut seeds were germinated (Day 1) and grown into seedlings (Day 10). Plant roots were treated with either nitrate ($KNO_3$) (Days 11 to 17) or mock-treated with water. On day 21, roots were inoculated with rhizobia (*Bradyrhizobium sp. SEMIA 6144*) after withholding water for 3 days. Again, plant roots were treated with either nitrate (Days 25 to 31) or mock plants continued with water treatment. Plant roots phenotype and tissue collection were performed at two time points, 7 days post-infection (DPI; Day 28) and 21 dpi (Day 42). Light and deep orange circles represent the no-watering period before rhizobia (Day 18 to 20) and after rhizobia (Day 22 to 24) treatment. Crossed watering icons also indicate time points without watering.

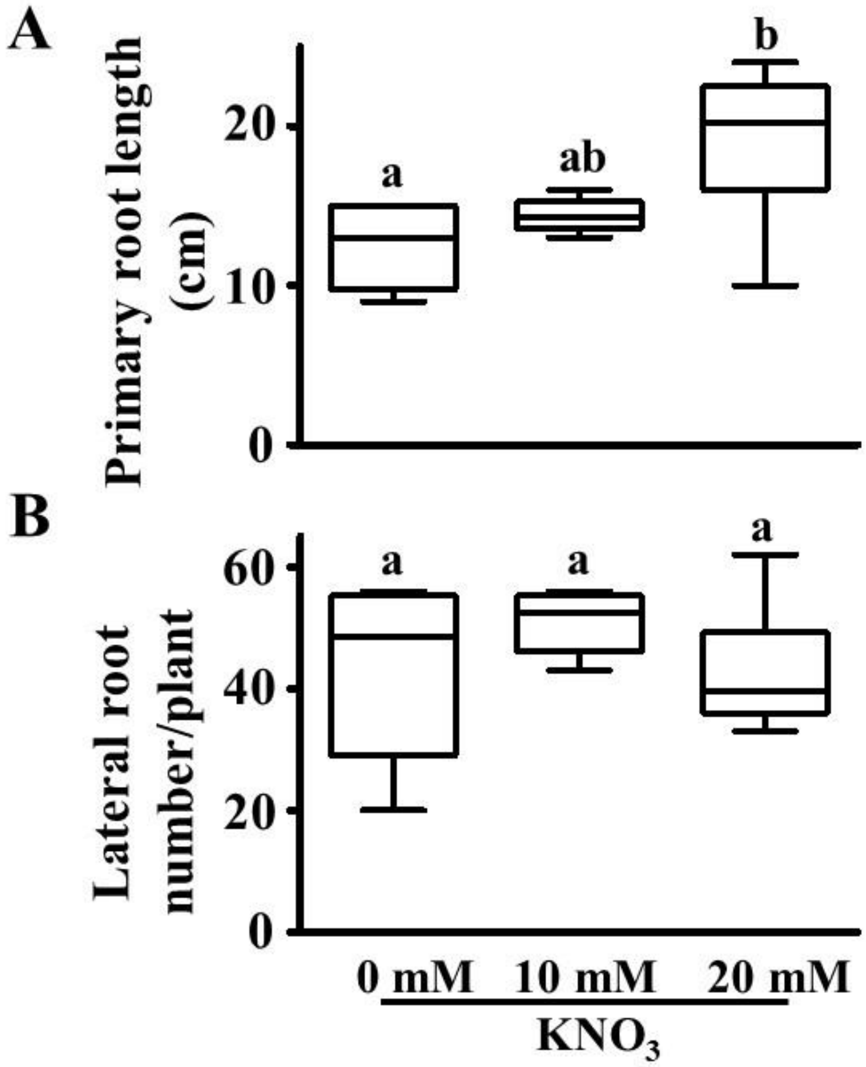


**Figure S2. Effect of different concentrations of nitrate on root architecture in peanut. A-B)** Peanut plants were treated with different concentrations of nitrate ($KNO_3$) and inoculated with rhizobia according to Figure S1. Subsequently, primary root length **(A)** and lateral root number **(B)** per plant at 21 dpi were quantified in rhizobia plus 10- or 20-mM nitrate-treated roots compared to only rhizobia-treated roots. Three biological replicates were used, and each replicate contained ~5 roots. Different letters indicate significant differences (Dunn's multiple-comparisons test, $P < 0.05$) between samples. Error bars represent ± standard error (SE).

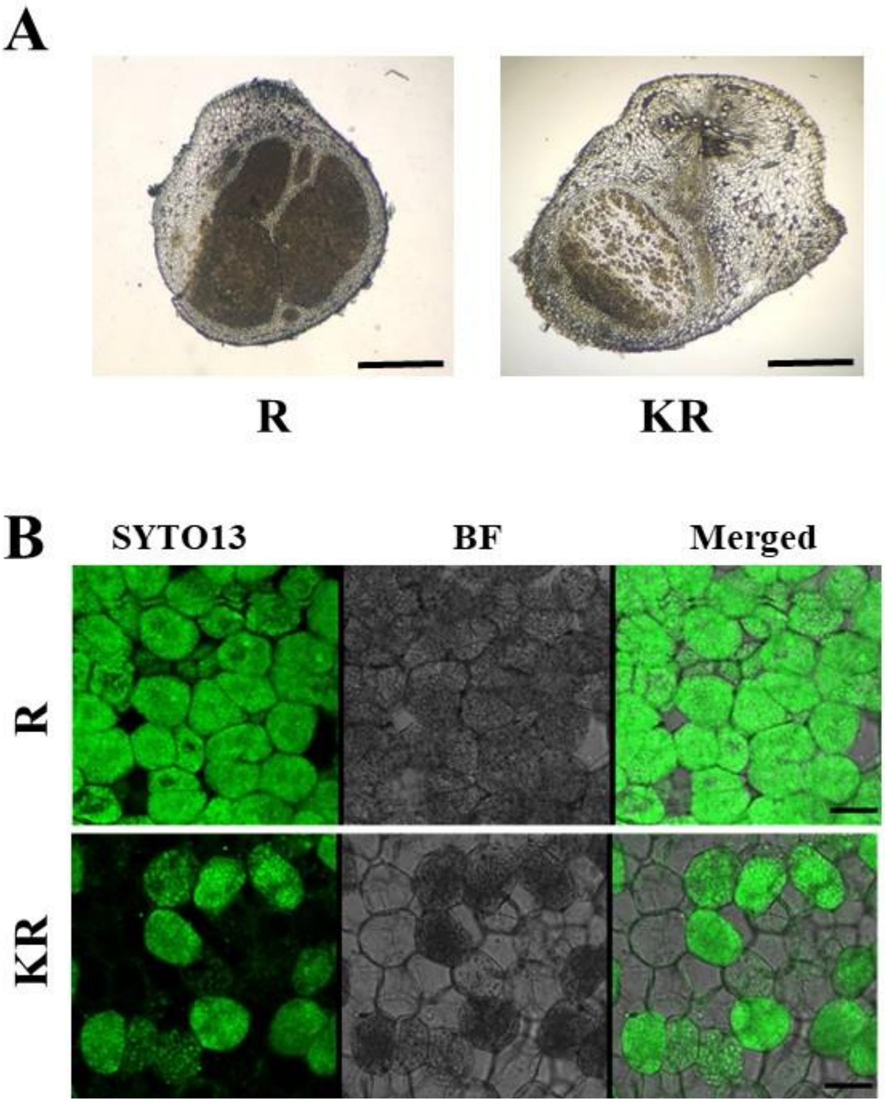


**Figure S3. Nitrate treatment restricts rhizobial colonization in mature nodules in peanut. A)** Light microscopy images of the transverse section (TS) of nodules (40 µm thickness), obtained using a vibratome and stained with toluidine blue, showed reduced rhizobial occupancy in rhizobia plus 20 mM nitrate-treated roots (KR) compared to rhizobia-treated roots (R). Scale bars = 0.5 mm. **B)** Confocal microscopy images of the transverse section (TS) of nodules (40 µm thickness), obtained using a vibratome and stained with SYTO13, exhibited reduced rhizobial occupancy in KR compared to R. Scale bars = 20 µm.

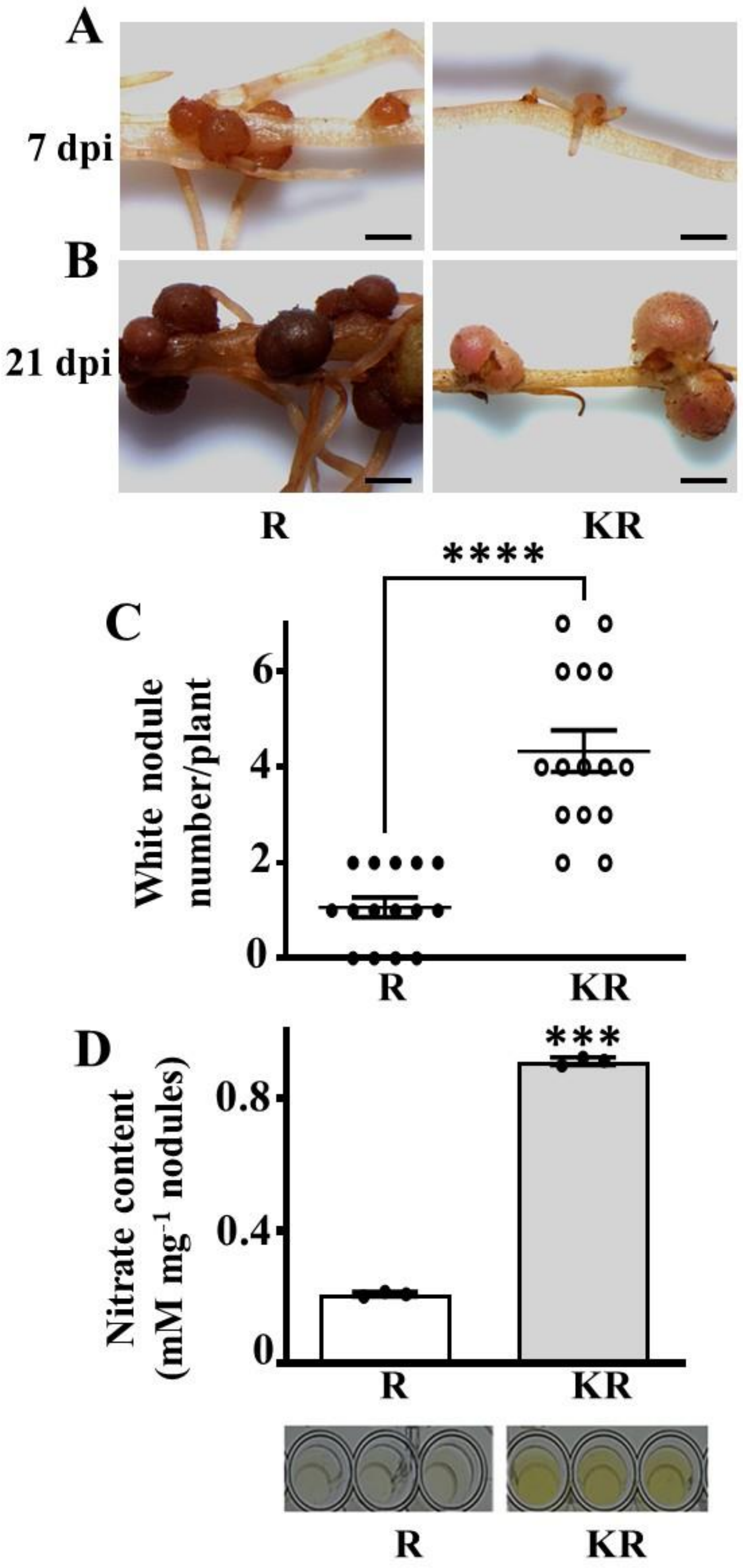
A
7 dpi
B
21 dpi
R
KR
****
C
White nodule number/plant
0
2
4
6
R
KR
D
***
Nitrate content (mM mg-1 nodules)
0
0.4
0.8
R
KR
R
KR

**Figure S4. Phenotype and biochemical features of nitrate-treated nodules in peanut. A and B)** Representative image of 0- (R or rhizobia) and 20 mM nitrate ($KNO_3$) (KR or nitrate plus rhizobia) treated roots at 7 dpi and 21 dpi. Scale bars = 2 mm. **C)** Quantification of white nodule numbers per hairy root in R and KR roots at 21 dpi. Three biological replicates were used, and each replicate contained 5 to 7 roots. Asterisks indicate statistically significant differences (Mann-Whitney U test, ****P < 0.0002) between samples. Error bars represent ±SE. **D)** Estimation of nitrate content in R and KR roots at 21 dpi. Three biological replicates were used for this assay. Asterisks indicate statistically significant differences (unpaired t test, P < 0.0001) between samples. Bars represent ± SE. A plate view of the reactions of the nitrate assay (lower panel).

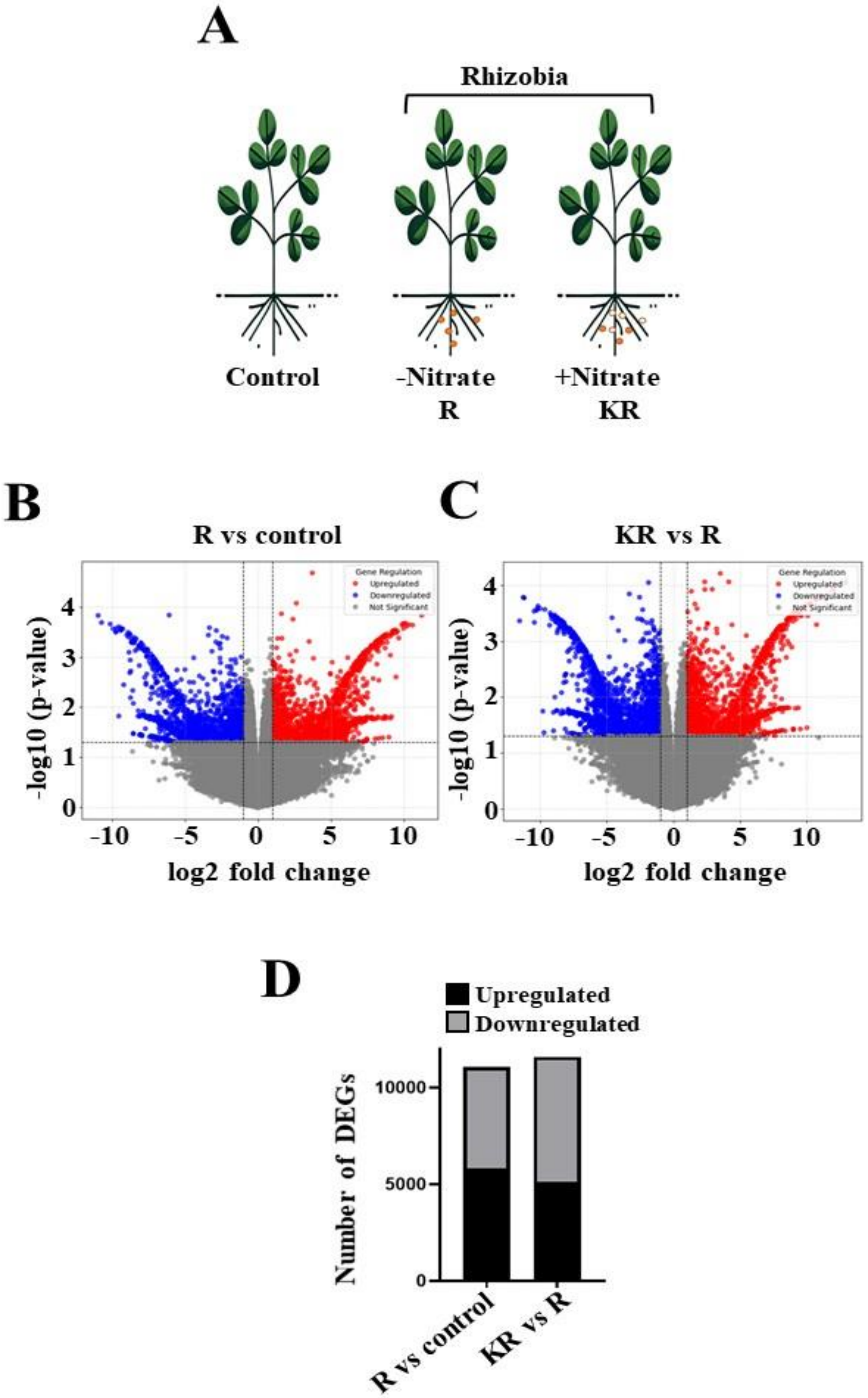
A
Rhizobia
Control
-Nitrate
R
+Nitrate
KR
B
R vs control
C
KR vs R
log2 fold change
-log10 (p-value)
D
Upregulated
Downregulated
Number of DEGs
R vs control
KR vs R

**Figure S5. Transcriptomic analysis of nitrate-treated roots. A)** A diagram showing the three types of root samples used for RNA-seq analysis. Ten-day-old peanut seedlings were treated with 20 mM nitrate ($KNO_3$) for one week and subsequently inoculated with rhizobia (KR). Ten-day-old peanut seedlings (mock) were treated with water for one week and then inoculated with rhizobia (R plants). For control, 10-day-old peanut seedlings were treated with water (mock) for 1 week, and not inoculated with rhizobia. Samples were collected at 6 dpi for RNA-seq analysis. **B and C)** Volcano plot illustrating the distribution of gene expression changes identified in peanut root RNA-seq data. The x-axis represents the log2 fold change, while the y-axis indicates the $-\log_{10}$ ($p$-value). Red dots correspond to significantly upregulated genes, blue dots represent significantly downregulated genes, and grey dots indicate non-significant changes. Vertical dashed lines denote the fold-change cut-off ($\log_2$ fold change, $\geq 1$), and the horizontal dashed line marks the significance threshold ($p < 0.05$). **D)** The number of differentially expressed genes (DEGs) were compared between R (Rhizobia) vs control roots, and KR vs R roots.

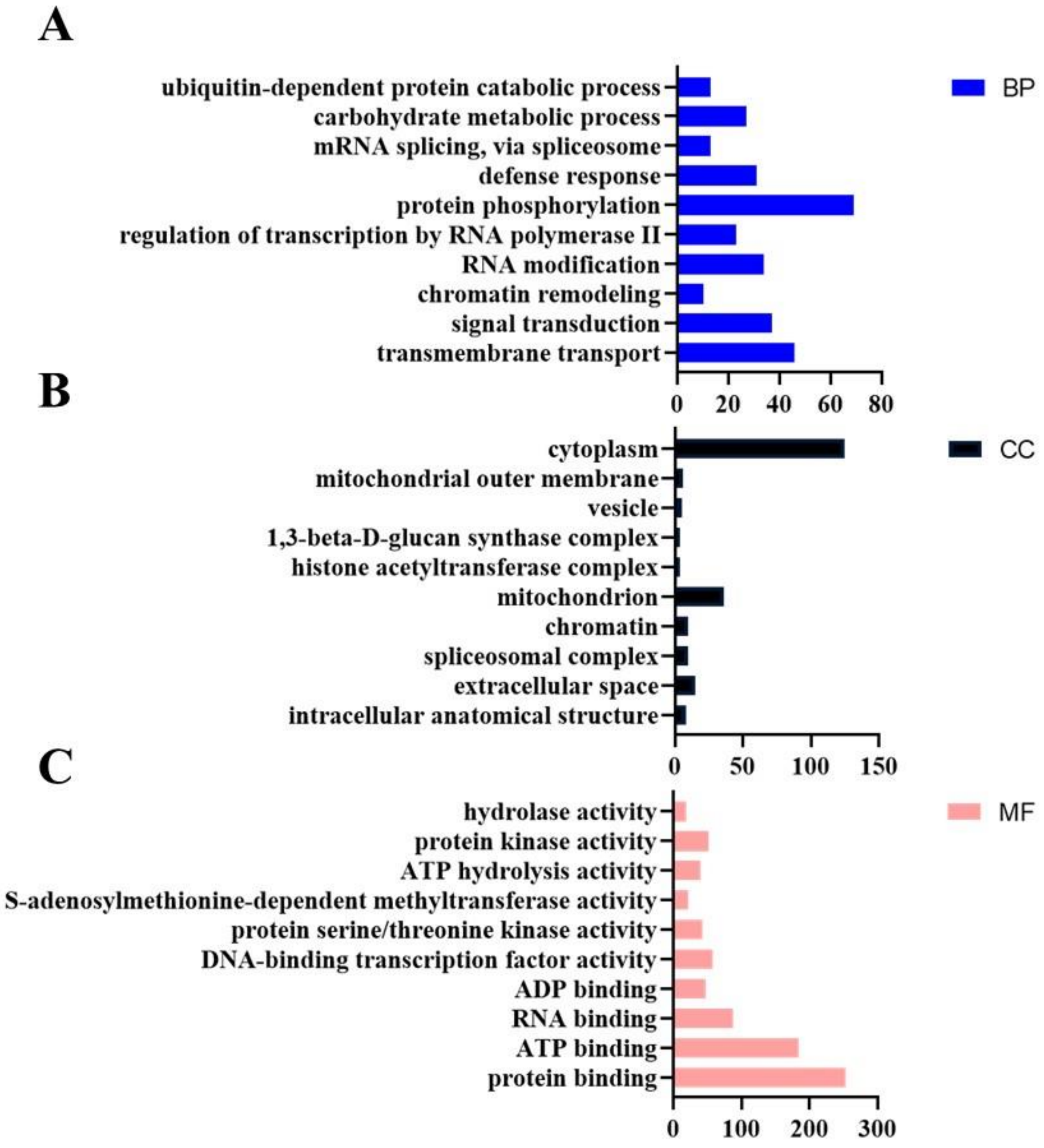


**Figure S6. GO analysis of common DEGs between upregulated genes in rhizobia-treated roots compared to control (R vs control) and downregulated genes in nitrate plus rhizobia-treated roots compared to rhizobia-treated roots (KR vs R).** The bar chart represents the top 10 enriched GO terms under the categories of biological process (BP), cellular component (CC) and molecular function (MF).

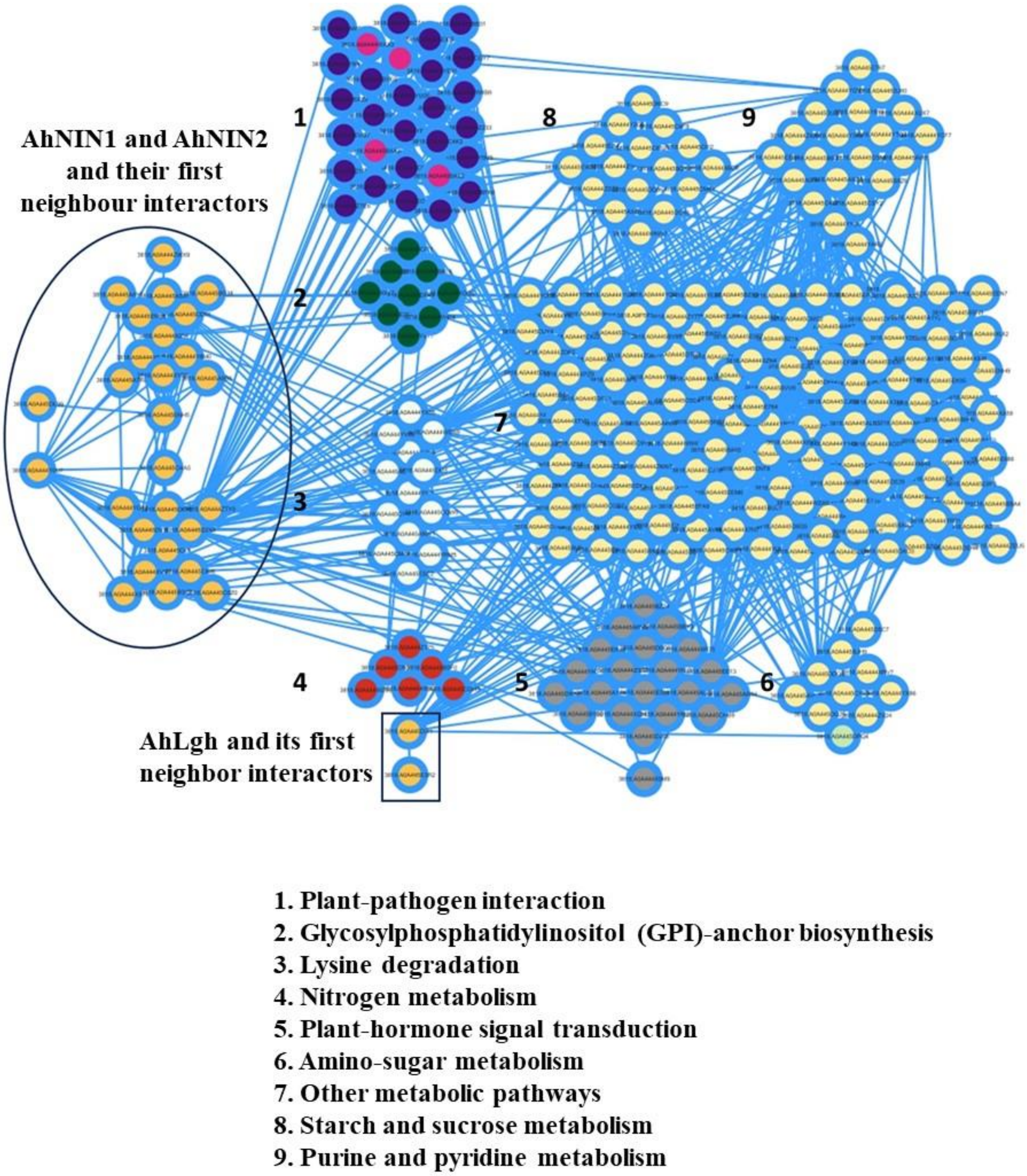


**Figure S7. Co-expression network of common DEGs between upregulated genes in rhizobia-treated roots compared to control (R vs control) and downregulated genes in nitrate plus rhizobia-treated roots compared to rhizobia-treated roots (KR vs R)**. This network highlights NIN interactions. Nine different functionally clustered modules were connected with AhNIN1 and AhNIN2. AhLgh is a part of one of the modules.

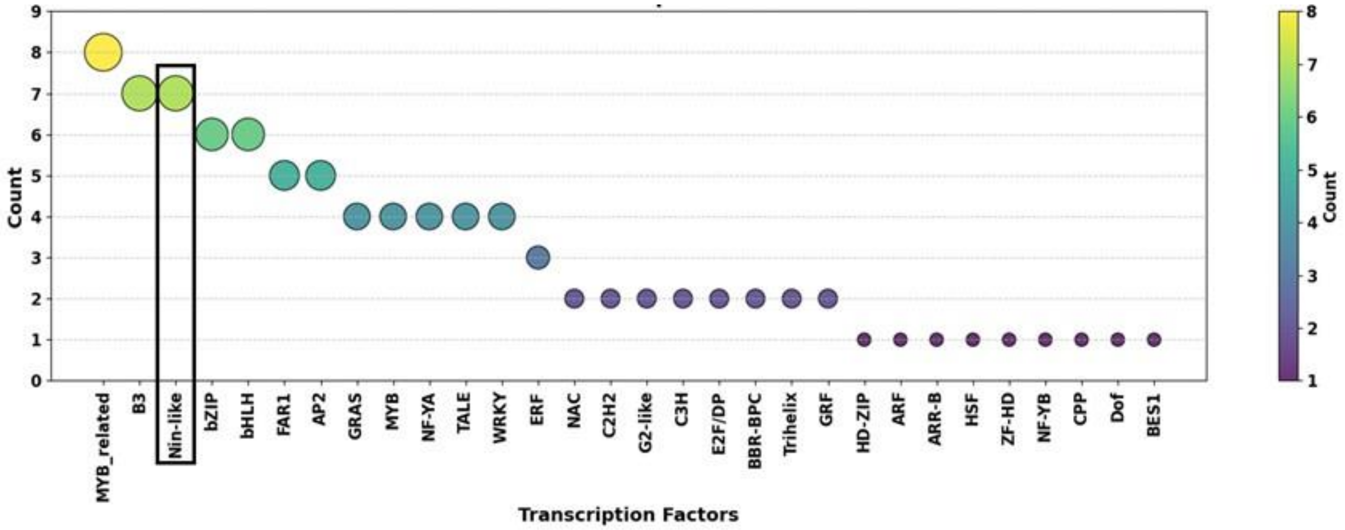


**Figure S8. Prediction of transcription factors (TFs) within common DEGs.** The dot plot represents the count of predicted TFs from the common DEGs between upregulated genes in rhizobia-treated roots compared to control (R vs control) and downregulated genes in nitrate plus rhizobia-treated roots compared to rhizobia-treated roots (KR vs R).

**A**

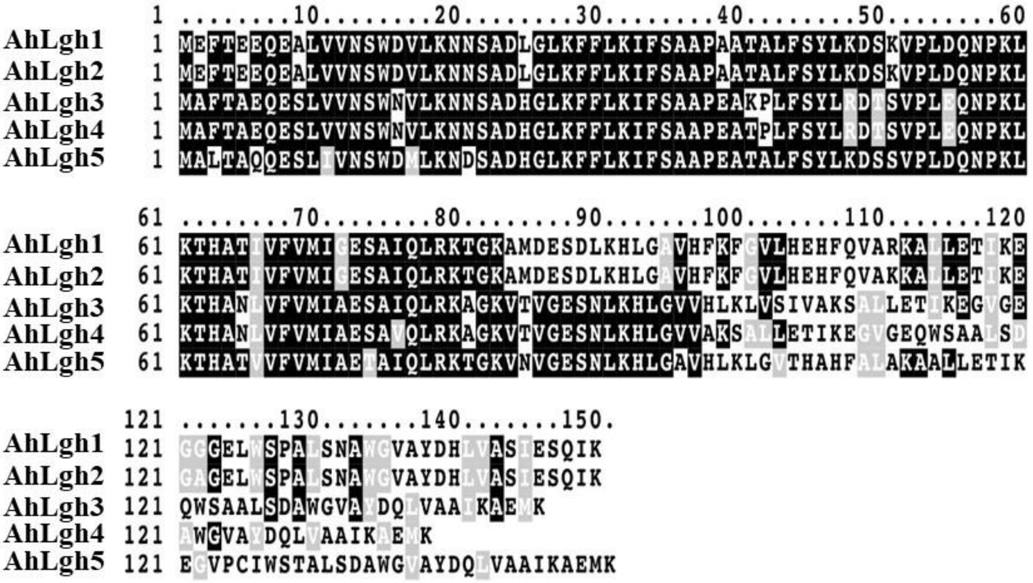


**B**

| | AhLgh1 | AhLgh2 | AhLgh3 | AhLgh4 | AhLgh5 |
|---|---|---|---|---|---|
| **AhLgh1** | 100 | 98.6667 | 69.1781 | 71.0145 | 72.6667 |
| **AhLgh2** | 98.6667 | 100 | 69.863 | 71.7391 | 73.3333 |
| **AhLgh3** | 69.1781 | 69.863 | 100 | 96.3768 | 80.137 |
| **AhLgh4** | 71.0145 | 71.7391 | 96.3768 | 100 | 81.8841 |
| **AhLgh5** | 72.6667 | 73.3333 | 80.137 | 81.8841 | 100 |

**Figure S9. Comparative sequence analysis of non-symbiotic leghemoglobins of peanut. A)** Multiple sequence alignment of five non-symbiotic leghemoglobin proteins represents a high degree of sequence similarity among them. **B)** Pairwise alignment score of sequence similarity among five non-symbiotic leghemoglobins.

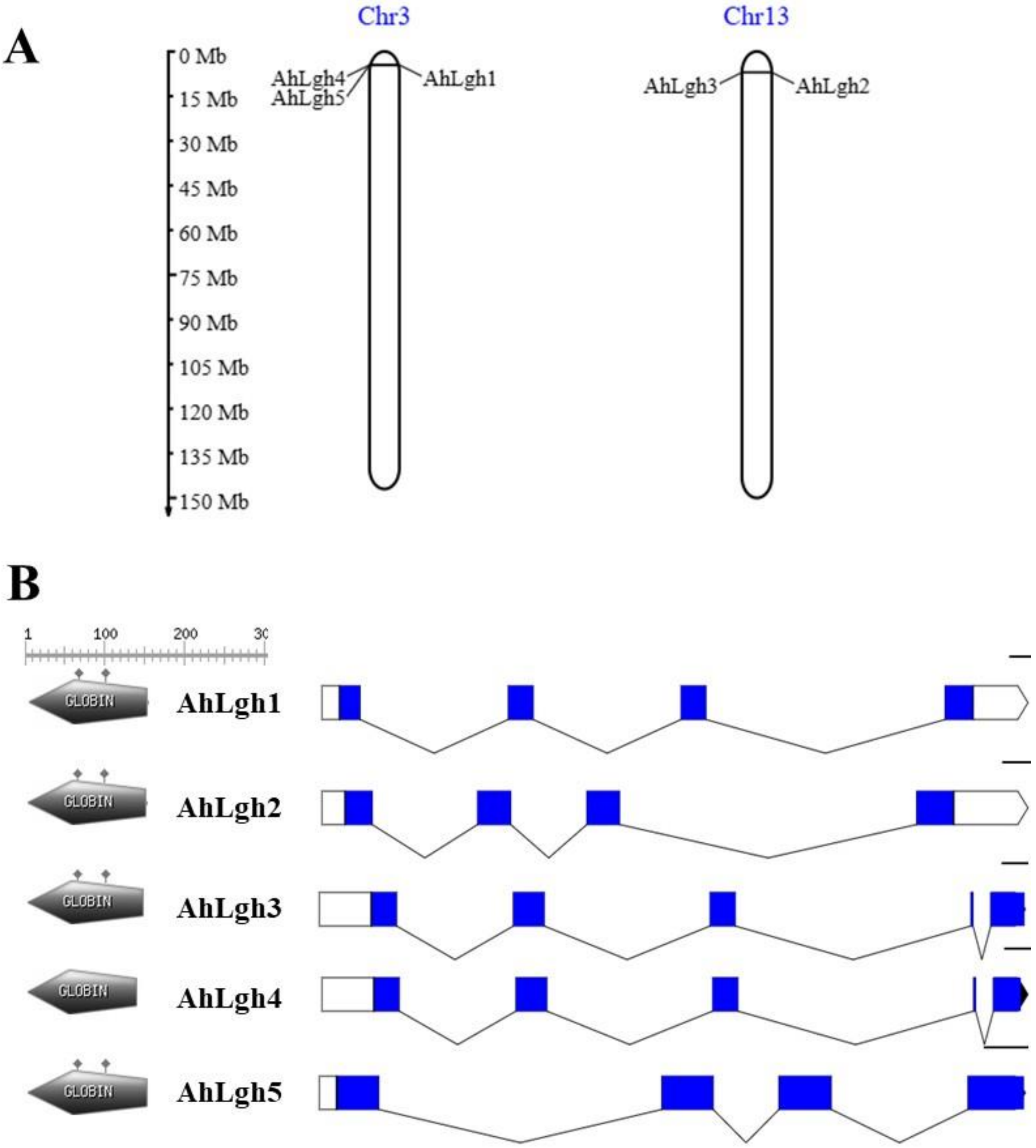


**Figure S10. Chromosomal localization and gene architecture of non-symbiotic leghemoglobin genes of peanut. A)** Chromosomal distribution pattern of five non-symbiotic leghemoglobin genes in peanut. The top values represent the number of chromosomes, and the scale is in megabase (Mb). **B)** The exon/intron organization of five non-symbiotic leghemoglobin genes. Blue boxes represent exons, black lines represent introns, and the white box denotes 5' UTR or 3' UTR.

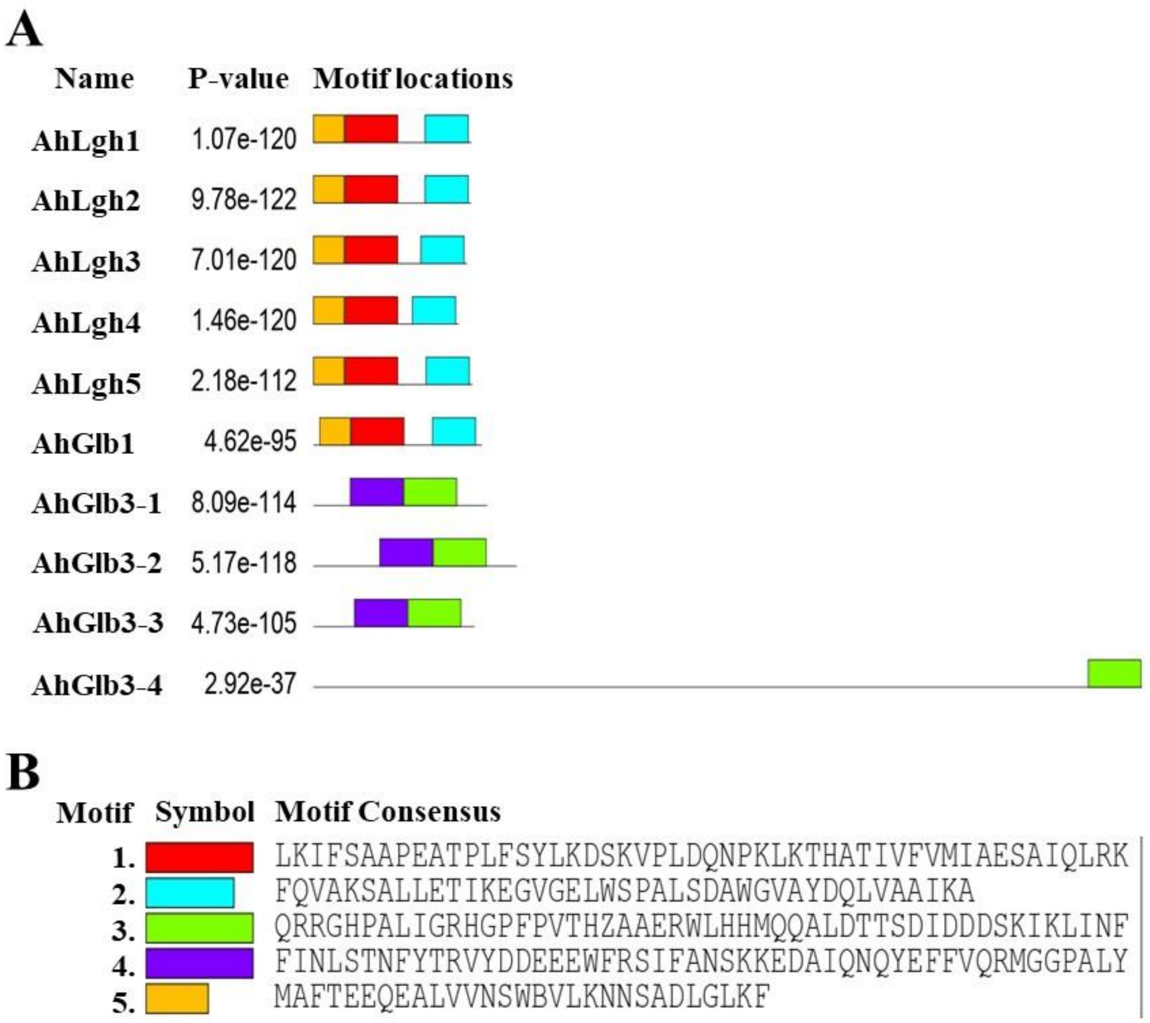


**Figure S11. Phylogenetic tree and conserved motif of leghemoglobins of peanut. A)** Distribution of conserved motifs of five non-symbiotic leghemoglobins and five Glbs in peanut. Different coloured boxes represent different motifs. **B)** Sequence logo of the conserved motifs.

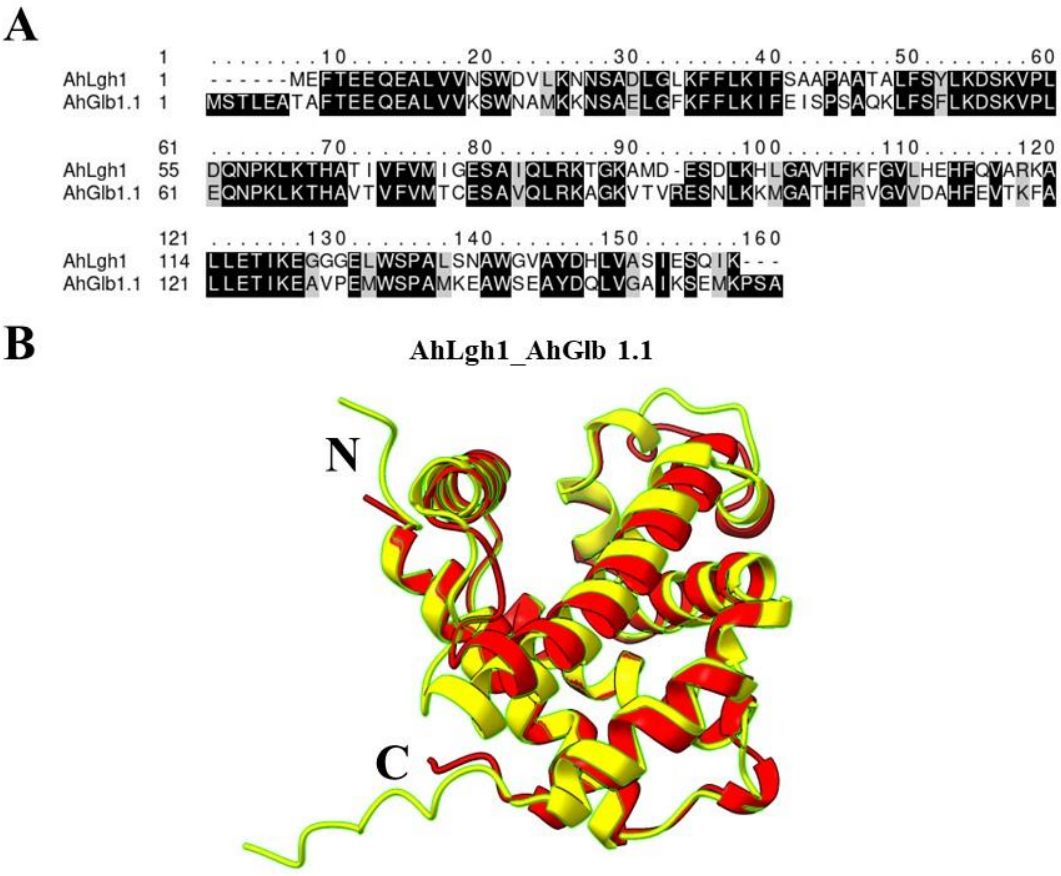


**Figure S12. Comparative sequence analysis of AhLgh1 with AhGlb1.1 of peanut. A)** Pairwise alignment of AhLgh1 and AhGlb1.1 proteins represents a lower degree of sequence similarity between them. **B)** Superimposition of cartoon models of AhLgh1 with AhGlb1.1 by aligned the 3D structure generated from AlphaFold3, using the MatchMaker tool in UCSF ChimeraX.

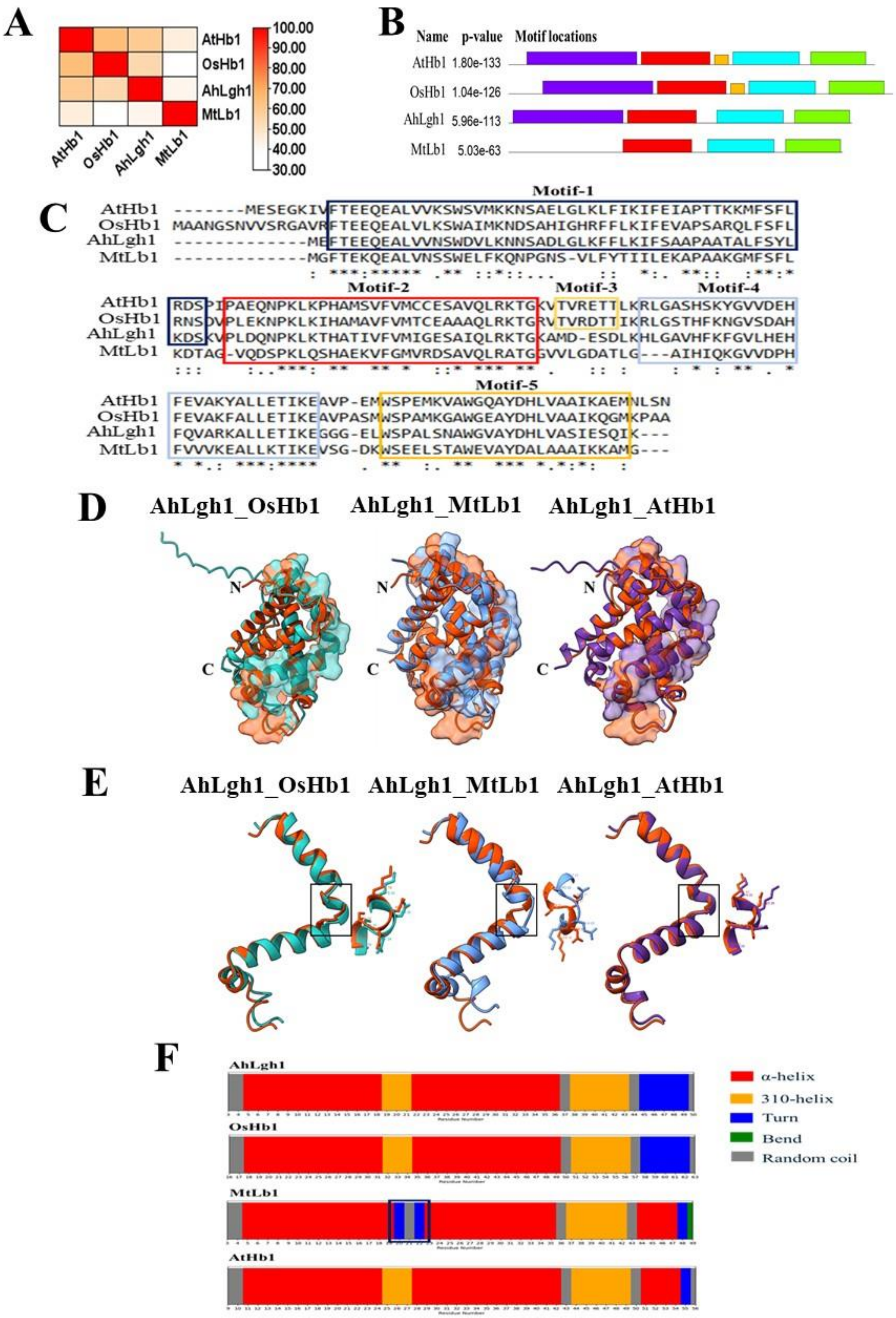
A
AtHb1
OsHb1
AhLgh1
MtLb1
100.00
90.00
80.00
70.00
60.00
50.00
40.00
30.00
B
Name p-value Motif locations
AtHb1 1.80e-133
OsHb1 1.04e-126
AhLgh1 5.96e-113
MtLb1 5.03e-63
C
Motif-1
AtHb1 -------MESEGKIVFTEEQEALVVKSWSVMKKNSAELGLKLFIKIFEIAPTTKKMFSFL
OsHb1 MAANGSNVVSRGAVRFTEEQEALVLKSWAIMKNDSAHIGHRFFLKIFEVAPSARQLFSFL
AhLgh1 -------------MEFTEEQEALVVNSWDVLKNNSADLGLKFFLKIFSAAPAATALFSYL
MtLb1 -------------MGFTEKQEALVNSSWELFKQNPGNS-VLFYTIILEKAPAAKGMFSFL
Motif-2 Motif-3 Motif-4
AtHb1 RDSPIPAEQNPKLKPHAMSVFVMCCESAVQLRKTGKVTVRETTLKRLGASHSKYGVVDEH
OsHb1 RNSDVPLEKNPKLKIHAMAVFVMTCEAAAQLRKTGRVTVRDTTIKRLGSTHFKNGVSDAH
AhLgh1 KDSKVPLDQNPKLKTHATIVFVMIGESAIQLRKTGKAMD-ESDLKHLGAVHFKFGVLHEH
MtLb1 KDTAG-VQDSPKLQSHAEKVFGMVRDSAVQLRATGGVVLGDATLG---AIHIQKGVVDPH
Motif-5
AtHb1 FEVAKYALLETIKEAVP-EMWSPEMKVAWGQAYDHLVAAIKAEMNLSN
OsHb1 FEVAKFALLETIKEAVPASMWSPAMKGAWGEAYDHLVAAIKQGMKPAA
AhLgh1 FQVARKALLETIKEGGG-ELWSPALSNAWGVAYDHLVASIESQIK---
MtLb1 FVVVKEALLKTIKEVSG-DKWSEELSTAWEVAYDALAAAIKKAMG---
D
AhLgh1_OsHb1
AhLgh1_MtLb1
AhLgh1_AtHb1
N
C
E
AhLgh1_OsHb1
AhLgh1_MtLb1
AhLgh1_AtHb1
F
AhLgh1
OsHb1
MtLb1
AtHb1
α-helix
310-helix
Turn
Bend
Random coil

**Figure S13. The structure of non-symbiotic leghemoglobin of peanut was compared with hemoglobins of legume and non-legumes**. **A)** A special group of non-symbiotic leghemoglobin protein sequences from *Arachis hypogaea* (AhLgh1) was compared with *Medicago truncatula* leghemoglobin (MtLb1) and other non-legume hemoglobins like *Arabidopsis thaliana* (AtHb1) and *Oryza sativa* (OsHb1). Multiple sequence alignment was performed using CLUSTALW (GenomeNet). Pairwise percentage similarities were calculated and visualised in the TB tool as a heatmap. Colour intensity reflects similarity scores, with darker shades indicating higher similarity. **B)** Protein sequences of AhLgh1, MtLb1, AtHb1 and OsHb1 were analyzed using MEME Suite for conserved motif identification. Each coloured box represents a distinct motif, with position and length proportional to their occurrence in the sequence. **C)** Multiple sequence alignment of AtHb1, OsHb1, AhLgh1 and MtLb1. All five conserved motifs identified by MEME were mapped into the sequence alignment and were highlighted with coloured boxes. The boxed regions correspond to functionally important domains, such as heme-binding or cofactor-binding residues. **D)** Predicted 3D structures of AhLgh1, OsHb1, MtLb1 and AtHb1 were generated in AlphaFold3 and aligned using the MatchMaker tool in UCSF ChimeraX. Superimposition of cartoon models of AhLgh1 with OsHb1, MtLb1 and AtHb1 illustrates the degree of structural conservation and divergence of *Arachis hypogaea* with legume (*Medicago truncatula*) and non-legume hemoglobins (*Arabidopsis thaliana* and *Oryza sativa*). AhLgh1 is orange red, OsHb1 is light sea green, MtLb1 is camphor blue and AtHb1 is Rebecca purple in colour. Motif 1 was highlighted in cartoon representation with a semi-transparent surface (60% transparency). **E)** Motif 1 regions of AhLgh1, OsHb1, MtLb1 and AtHb1 were mined from the predicted 3D superimposed structures. Pairwise alignments highlighted structural similarities and differences of AhLgh1–OsHb1, AhLgh1–MtLb1 and AhLgh1–AtHb1 specifically within motif 1. The right side is the zoom view of the region marked inside the black box. **F)** Secondary structures of motif 1 were generated using DSSP using GROMACS package and python for AhLgh1, OsHb1, MtLb1 and AtHb1. The figure highlights α-helix, 310-helix, turn, bends and random coil regions within motif 1, showing conserved and divergent structural features of AhLgh with legume and non-legume hemoglobins.

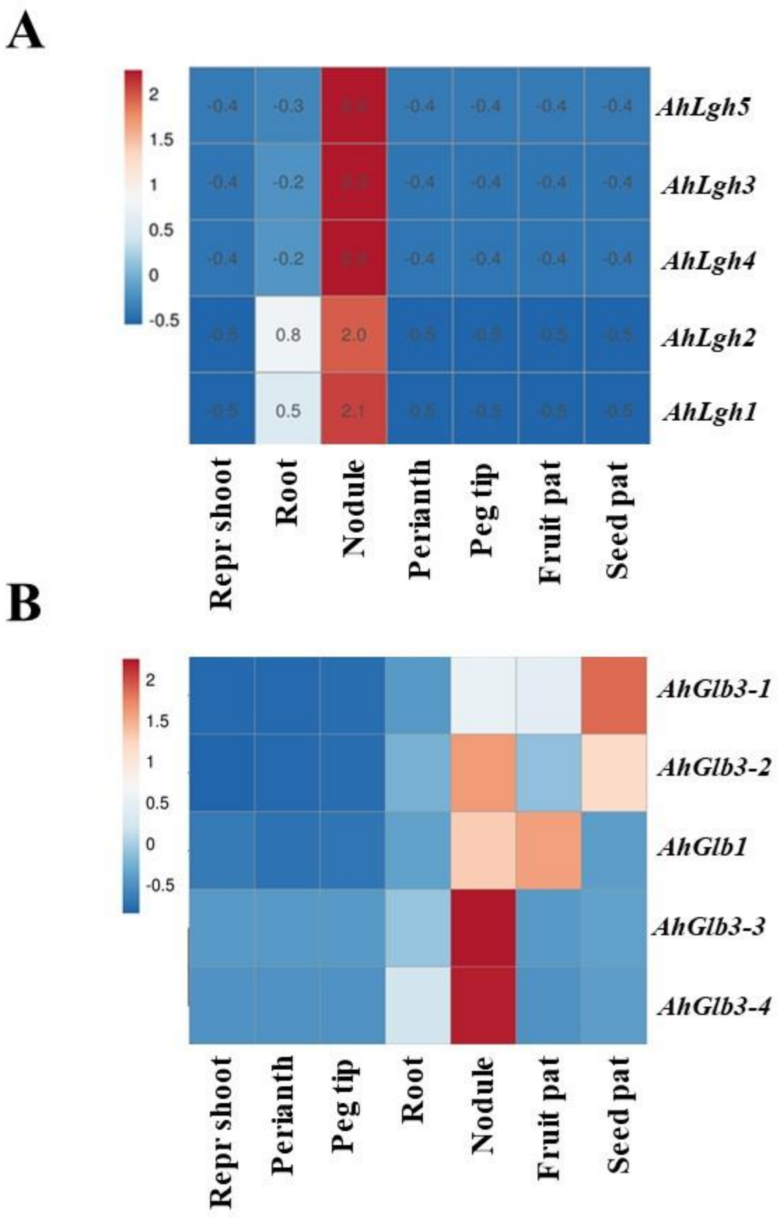


**Figure S14. Expression profiles of non-symbiotic leghemoglobin genes in peanut. A)** Tissue-specific expression of five non-symbiotic leghemoglobin genes of peanut in different vegetative and reproductive tissues. Data was retrieved from the peanut database (https://www.peanutbase.org/). **B)** Tissue-specific expression of five Glbs genes of peanut in different vegetative and reproductive tissues. Data was retrieved from the peanut database (https://www.peanutbase.org/).

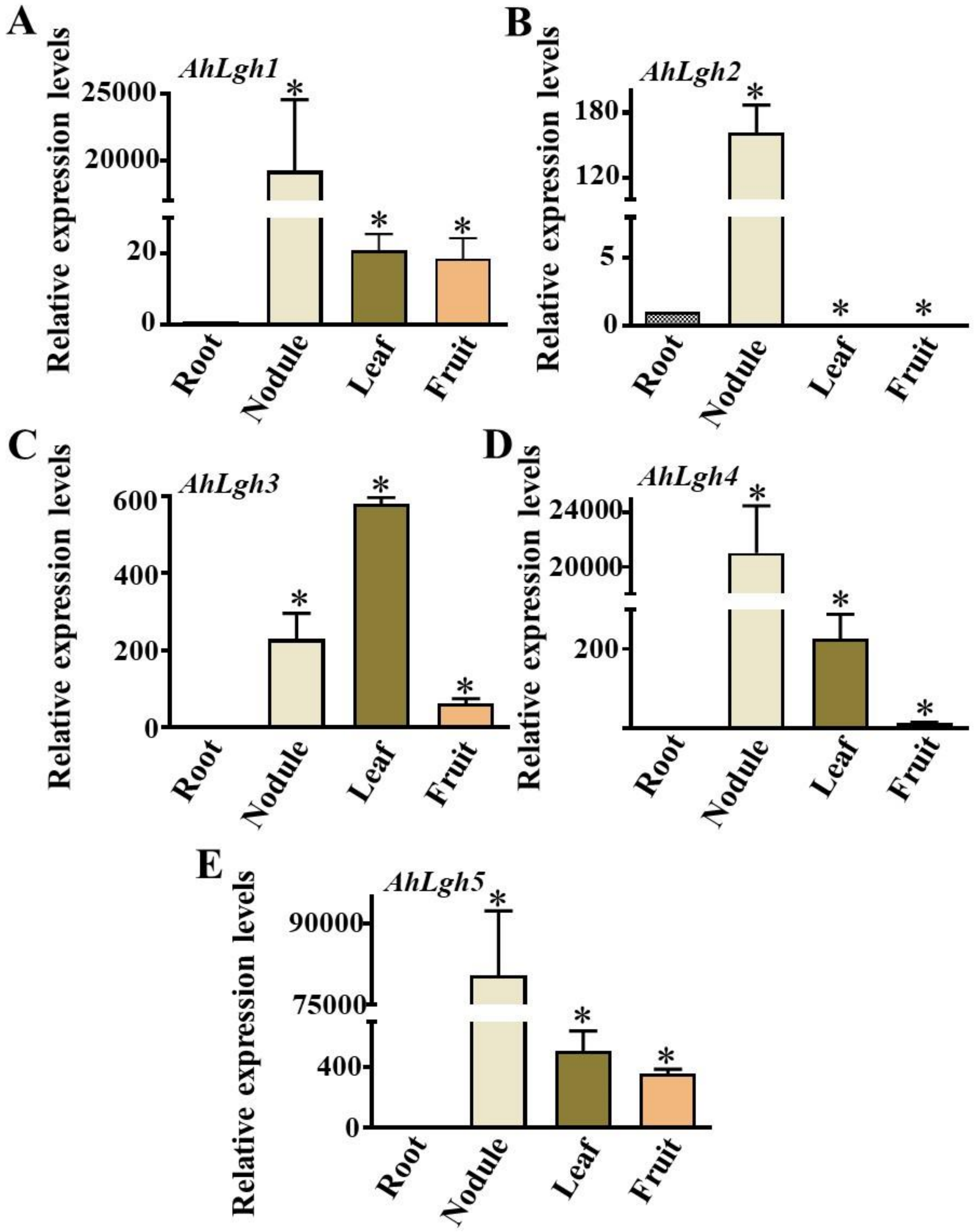


**Figure S15. Tissue-specific expression of non-symbiotic leghemoglobin genes in peanut. A to E)** Tissue-specific expression (Root, Nodule, Leaf and Fruit) of *AhLgh1*, *AhLgh2, AhLgh3*, *AhLgh4 and AhLgh5* in peanut (*Arachis hypogaea*) was measured by RT-qPCR. The gene expression levels were normalized using *AhActin*. Expression in roots was set at 1. Three biological replicates were used. Error bars represent ±SE. Asterisks indicate statistically significant differences (Student's t-test, *P < 0.05) between samples.

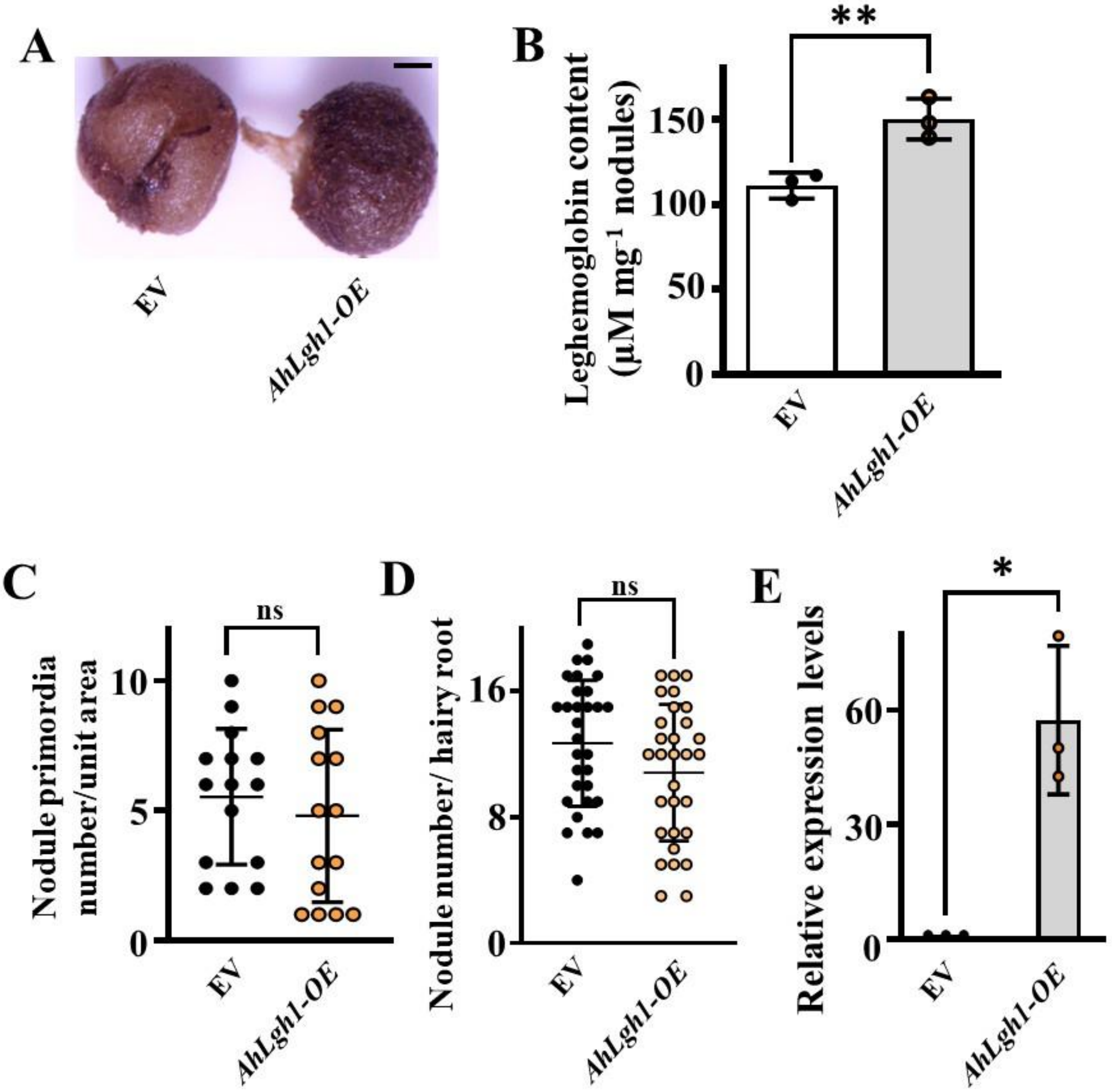
A
EV
AhLgh1-OE
B
**
Leghemoglobin content
(μM mg-1 nodules)
150
100
50
0
EV
AhLgh1-OE
C
ns
Nodule primordia
number/unit area
10
5
0
EV
AhLgh1-OE
D
ns
Nodule number/ hairy root
16
8
0
EV
AhLgh1-OE
E
*
Relative expression levels
60
30
0
EV
AhLgh1-OE

**Figure S16. Overexpression of *AhLgh1* increases leghemogobin content. A)** Representative light microscopy images exhibiting nodule colour differences of hairy roots transformed with EV and *OE-AhLgh1*. Scale bars = 0.5 mm. **B)** Estimation of leghemoglobin content in nodules of empty vector (EV) and *OE-AhLgh1* transformed hairy roots at 21 dpi. Asterisks indicate statistically significant differences (unpaired t test, **P < 0.009) between samples. Error bars represent ±SE. **C)** Nodule primordia numbers were counted at 7 dpi in empty vector (EV) and *OE-AhLgh1* transformed hairy root plants inoculated with *Bradyrhizobium sp. SEMIA 6144*. **D)** Nodule numbers per hairy roots were counted at 21 dpi in empty vector (EV) and *OE-AhLgh1* transformed hairy root plants inoculated with *Bradyrhizobium sp. SEMIA 6144.* Three biological replicates were used, and each replicate contained 5 to 7 roots and 10 to 12 plants for the estimation of nodule primordia and nodule number, respectively. Asterisks indicate statistically significant differences (Mann-Whitney U test) between samples. Error bars represent ±SE. **E)** Transcript levels of *AhLgh1* were detected by RT-qPCR in transgenic hairy roots of EV and *OE-AhLgh1* lines at 21 dpi by RT-qPCR. The expression values were normalized against the peanut *Actin* gene expression. Expression in EV roots was set at 1. Three biological replicates were used. Error bars represent ±SE. Asterisks indicate statistically significant differences (Student's t-test, *P < 0.01) between samples.

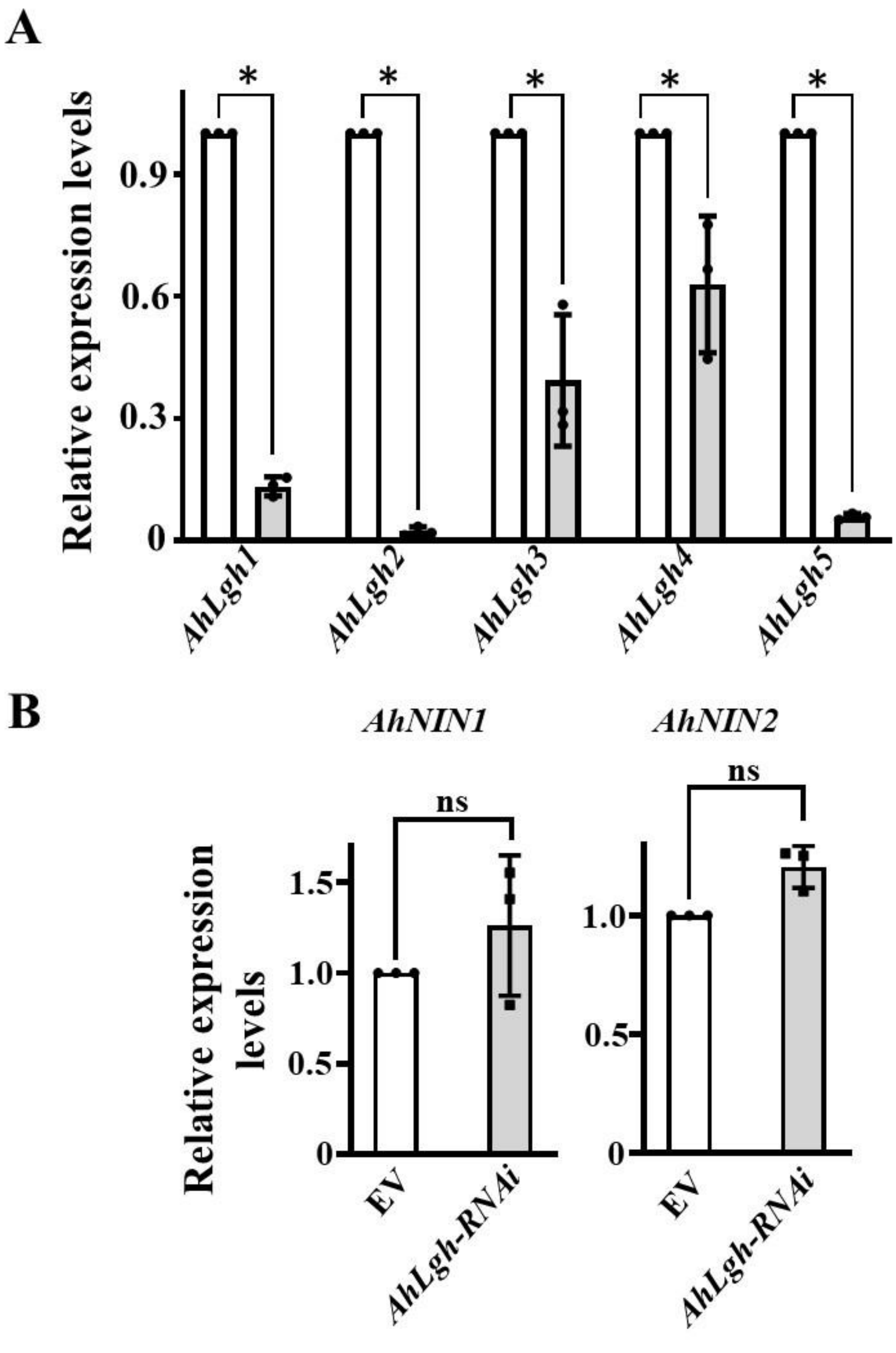


**Figure S17. NIN acts upstream of *AhLgh1*. A)** Transcript levels of *AhLgh1*, *AhLgh2, AhLgh3*, *AhLgh4* and *AhLgh5* were detected by RT-qPCR in transgenic hairy roots of EV and *AhNIN-RNAi* lines at 21 dpi. **B)** Transcript levels of *AhNIN1* and *AhNIN2* were detected by RT-qPCR in transgenic hairy roots of EV and *AhLgh-RNAi* lines at 21 dpi. The gene expression levels were normalized using *AhActin*. For RT-qPCR, three biological replicates were used. Bars represent ± SE. Asterisks indicate statistically significant differences (Student's t-test, *P < 0.01) between samples.

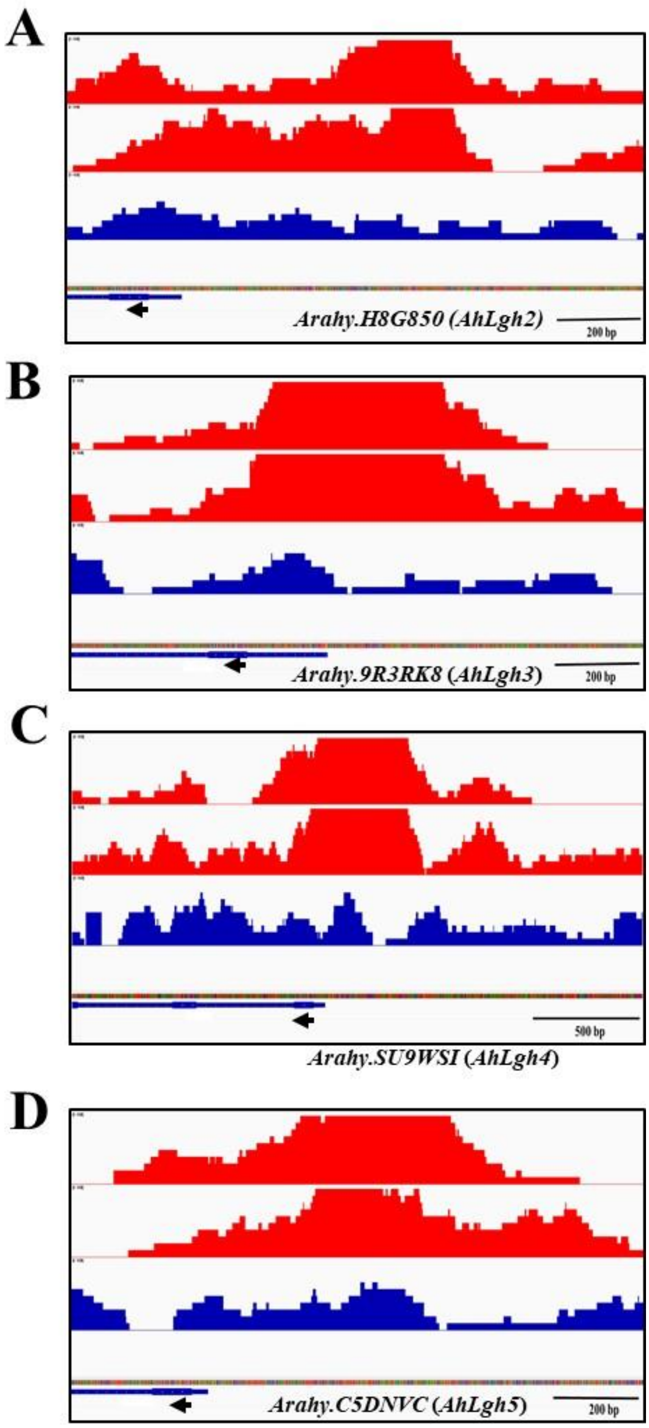


**Figure S18. NIN binds to the NRCE of AhLgh's promoter region. A to D)** Localization of the enriched binding peaks of AhNIN in the promoter regions of non-symbiotic leghemoglobin genes; *AhLgh2*, *AhLgh3*, *AhLgh4* and *AhLgh5*. The arrow indicates the translation start site and its orientation. Two replicates were used. Input: sequencing results of genomic DNA fragments used as a negative control.

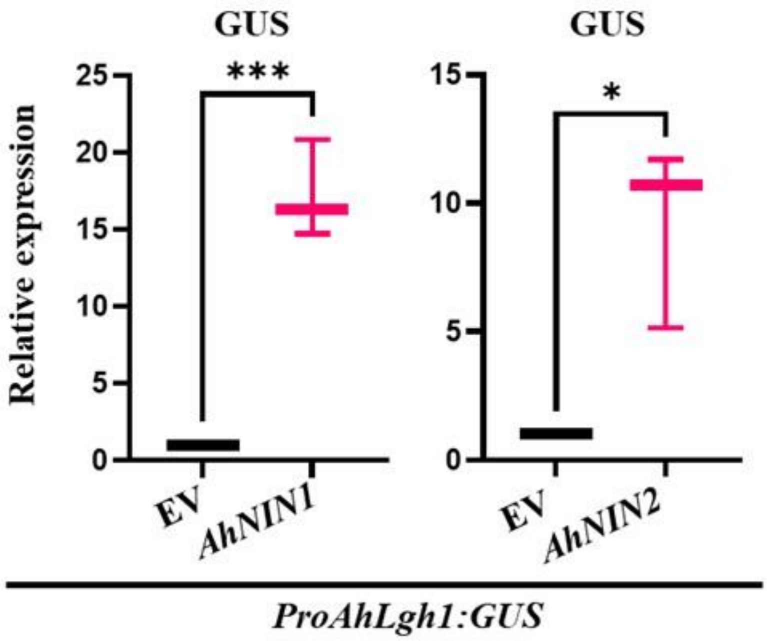


**Figure S19. Quantification of *GUS* gene expression. A to B)** Relative expression of *β-glucuronidase* (*GUS*) was quantified in *N. benthamiana* leaves co-transformed with EV, *proAhLgh1:GUS* or *AhNIN's (AhNIN1 and AhNIN2)* and *proAhLgh1:GUS.* The gene expression levels were normalized using *AhActin*. For RT-qPCR, three biological replicates were used. Bars represent ± SE. Asterisks indicate statistically significant differences (Student's t-test, *P < 0.01 and, ***P < 0.001) between samples.
