## Supplementary material for "Nitrate restricts the expression of non-symbiotic leghemoglobin through inhibition of nodule inception protein in nodules of peanut (*Arachis hypogaea*)"

### Transcriptome analysis

Differential gene expression analysis was performed using the limma-voom package within the Galaxy platform (https://usegalaxy.org/). Rhizobia-treated (R) samples were compared to the control samples (R vs control) and rhizobia plus nitrate-treated (KR) samples were compared to the Rhizobia-treated samples (Kr vs R). Differential expression analysis was performed with an adjusted p-value threshold of 0.05 or less. For volcano plots, transcripts with log2 fold change equal or greater than 1 were marked as upregulated and those with log2 fold change equal or less than −1 as downregulated. To restrict the DEG set for functional analysis, we applied a more stringent cut-off of log2 fold change, ≥ 2 as upregulated and ≤ −2 as downregulated. Common genes between datasets were identified using the Venn diagram tool Venny 2.1 (Oliveros, 2007). The DEGs were plotted with log2 fold change values in heatmaps using TBtools (Chen et al., 2020). Functional annotation of assembled transcripts was performed by mapping GI identifiers to Gene Ontology (GO) terms and Kyoto Encyclopedia of Genes and Genomes (KEGG) pathways using the DAVID Functional Annotation Bioinformatics Microarray Analysis platform (https://davidbioinformatics.nih.gov/home.jsp) (Huang et al., 2009; Sherman et al., 2022). Transcription factor (TF) prediction was performed using the Plant Transcription Factor Database (https://planttfdb.gao-lab.org/) (Jin et al., 2017). To gain insights into the functional associations among combinations of nitrate and rhizobia-responsive proteins, a co-expression network was constructed using STRING version: 12.0 (Szklarczyk et al., 2023) and visualized in Cytoscape version 3.10.3 (Shannon et al., 2003). Using KEGG pathway enrichment, the network was subdivided into nine distinct functional clusters, each visualized with a unique colour (Metabolic pathways that fall into more than one clusters are painted with the same colour).

### Protein structure modeling, alignment, and DSSP analysis

The three-dimensional structures of AhLgh1, AhGlb1, MtLb1, OsHb1, and AtHb1, were generated using AlphaFold3 (https://alphafoldserver.com/) based on their amino acid sequences retrieved from phytozome database (https://phytozome-next.jgi.doe.gov/) (Abramson et al., 2024). All the predicted structures were validated by ERRAT and PROCHECK using SAVES v6.1 (https://saves.mbi.ucla.edu/) (Colovos and Yeates, 1993). Structural similarity was evaluated in UCSF ChimeraX using the MatchMaker tool (Pettersen et al., 2021). Alignment scores and root mean square deviation (RMSD) values were calculated to determine the relative similarity of AhLgh1 with AhGlb1, MtLb1, OsHb1, and AtHb1, respectively. Secondary structure elements were assigned with the Define Secondary Structure of Proteins (DSSP) algorithm as implemented in GROMACS (Abraham et al., 2015). The mkdssp module was applied to each model, and outputs were processed using custom Python scripts to calculate secondary structure composition.

**Nitrate content estimation**

Extraction was carried out with 10 volumes of sterilized Milli-Q water before incubation at 100°C for 20 minutes. After centrifugation, collected supernatant was added to Reaction Reagent 1 ($H_2SO_4$ + salicylic acid) and then Reaction Reagent 2 (8% NaOH) according to Hachiya and Okamoto, 2017. Finally, absorbance was detected by a spectrophotometer at 410 nm.

**Evolutionary analysis**

In order to generate the phylogenetic tree, protein sequences of non-symbiotic leghemoglobin genes from *Arachis hypogaea* were collected from the Phytozome database (*Arachis hypogaea* v1.0) and PeanutBase (https://www.peanutbase.org/). The CLUSTALW with default parameters were used to prepare multiple sequence alignment, and subsequently, IQ-TREE with the bootstrap method of 1000 replicates was used to obtain a reliable tree (Trifinopoulos et al., 2016). The exon-intron structures were visualized using the Exon-Intron Graphic Maker (http://www.wormweb.org/), where genomic sequences were aligned with CDS to distinguish exon and intron regions. The chromosomal distribution was visualized using MapGene2Chrom (MG2C) tool (http://mg2c.iask.in/mg2c_v1.0) (Jiangtao et al., 2015). The conserved motifs were detected by MEME Suite (https://meme-suite.org/meme/tools/meme), with the maximum number of motifs set to 5 for identification (Bailey and Elkan, 1994; Bailey et al., 2006).
